# Globally abundant birds disproportionately inhabit anthropogenic environments

**DOI:** 10.1101/2023.12.11.571069

**Authors:** Tadhg Carroll, Jack H. Hatfield, Chris D. Thomas

**Affiliations:** Leverhulme Centre for Anthropocene Biodiversity, University of York; Wentworth Way, York YO10 5DD, UK; Department of Biology, University of York; Wentworth Way, York YO10 5DD, UK

**Author notes:** These authors contributed equally to this work (Tadhg Carroll, Jack H. Hatfield).

## Abstract

Research into biodiversity change predominantly focuses on rarity and declines, but many ecological processes are governed by abundant species. Analysing 3,146 terrestrial bird species across 5,454 field-sampled sites, we find that three times more species in the top quartile for global abundance are more likely to occur in ecosystems characterised by major human land-cover modification (croplands, plantation forest, urban areas and pasture), compared with species in the bottom quartile. The likelihood of inhabiting human-modified environments consistently increases with global abundance across species with different dietary requirements, whereas low abundance species tend to have increased probabilities of occurrence (within their ranges) in relatively unmodified environments. Our findings suggest that human modification of the Earth’s land surface has favoured ‘anthrophilic’ species able to thrive in widespread anthromes.

**One-Sentence Summary:** The world’s most abundant birds disproportionately inhabit croplands, plantation forests, urban areas and pastures.

## Main Text

Human-associated land use has shaped the Earth for thousands of years (*1, 2*), appropriating and modifying large fractions of the planet’s terrestrial ice-free land and primary productivity. This has resulted in concerns over pervasive biodiversity declines (*3*), including falling species richness and population densities, and fears that a small subset of cosmopolitan species is prospering (and becoming locally abundant (*4*)) at the expense of other taxa (*5*). Rather than being a small subset, however, we find here that very large numbers of bird species with large global populations are associated with human-modified ecosystems, or anthromes. Half of the top quarter, by global abundance, of the world’s terrestrial bird species are now disproportionately associated with anthromes, rather than with more natural ecosystem types.

To become globally abundant, species require at least one of a number of characteristics (*6*–*8*). They must either have a wide niche breadth able to occupy a variety of environments (*9*) and/or be sufficiently well adapted to be numerous in a particularly widespread environments (*10, 11*). Smaller sized species, species using energy-rich food and those in particular climatic regions may also achieve high global abundance via high densities (*12*). As such, there are a number of reasons we might expect anthromes to be filled with globally abundant species.

Firstly, wide niche breadth may provide resilience in the face of anthropogenic change. Secondly, the majority of global landscapes contain at least some human-modified land (*1, 13*). These anthromes – particularly the croplands, pastures, plantations and urban areas considered here – are often structurally and biologically (e.g., the same crop species) similar across regions and continents (*14*), meaning that species able to use these environments may have the opportunity to assume geographically extensive distributions (*15*), sometimes aided by human introductions (*16*). Lastly, agricultural anthromes are geared toward high levels of production of certain crops and pasture plants, while urban environments may contain discarded ‘waste’ that represents a resource for some species (*17*). This could lead to large total abundances for species able to exploit human-associated resources.

Here, we investigate the relationship between the global abundance of bird species and their occurrence across a range of land-cover types which vary in intensity of contemporary human activity from ‘primary’ vegetation (least modified) to urban environments. Birds are especially suitable for this analysis because of their global distribution, the wide array of habitat and dietary niches they occupy, the key functional roles they play (*18*), and the availability of suitable data. Combining recent global abundance estimates for the world’s avifauna (*19*) with a global dataset of species occurrences in more and less human modified land cover types (*20, 21*) enabled us to examine whether the world’s more abundant birds are also those predominantly associated with anthromes, and whether any associations revealed are consistent across species with different dietary needs (‘trophic niches’ (*22*)). Our analysis included 5,454 individual survey sites (from the PREDICTS database (*20, 21*)) across all continents (except Antarctica; **Fig. 1**) and 3,146 species (almost a third of all known bird species), with good proportional representation across the entire distribution of species global abundances (**figs. S1-S2**), and of species inhabiting differing biogeographic realms, trophic niches and primary habitat usage, relative to the global avifauna (**Tables S1-S3** (*22, 23*)).

**Fig. 1.**
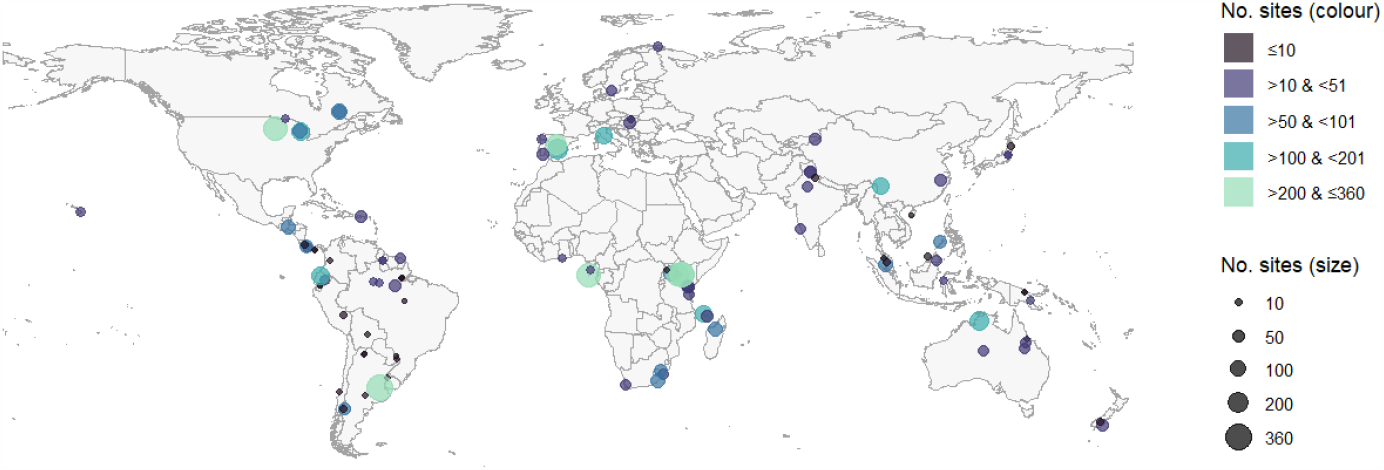
Location and number of sites for each PREDICTS study used in the analysis. Size of points varies continuously with the number of sites within studies. Colour of points represents discrete bins of the same quantity as a further visual aid.

Results from hierarchical logistic regression analyses reveal a clear progression of diverging trends in occurrence probability vs species’ global abundance (***Fig. 2***). The presence of the world’s more abundant terrestrial bird species is disproportionately associated with anthromes (i.e., cropland, pasture, plantation forest and urban areas) and disturbed ecosystems (young secondary vegetation; which often arises from recent human activities such as harvesting, fire and land abandonment), while less abundant species are disproportionately associated with the least anthromic vegetation types (primary, mature secondary and intermediate aged secondary vegetation). Effect sizes are greater (i.e., steeper slopes) and more certain for anthromic land-cover types (all 95% Credible Intervals (CRIs) are fully positive), whereas slope estimates for primary and secondary vegetation types are shallower and have 95% CRIs overlapping zero (***fig. S3*** and ***Table S4***). Note that all estimates of occurrence are for regions where a species was present (i.e., it was recorded within a particular study), and a given land cover type was surveyed in that study (i.e., the land cover type was ‘available’ to be occupied).

**Fig. 2.**
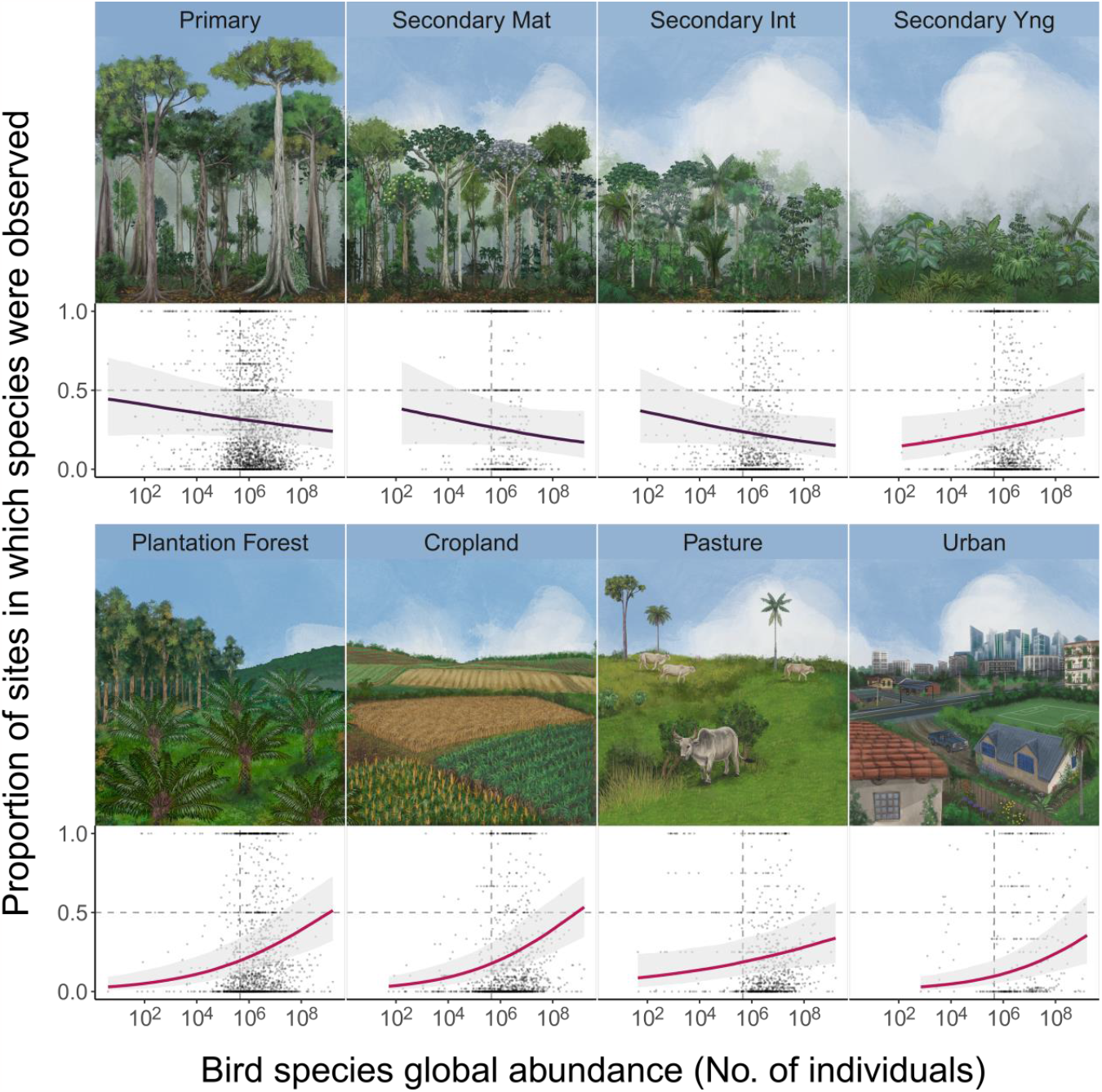
Diverging trends in occurrence probability vs bird species global abundance between more and less human modified land-cover types. X-axes represent species’ global abundance estimates (*19*) and Y-axes show proportions of land-cover specific survey sites in which bird species were recorded. Points in each panel show empirical, species-specific proportions of sites occupied in a given land-cover type, for each species recorded in at least one PREDICTS (*20*) study surveying which recorded that species. For secondary vegetation, Mat = Mature, Int = Intermediate aged, Yng = Young. Points falling along horizontal dashed lines indicate that a species was recorded in 50% of land-cover specific survey sites. Vertical dashed lines indicate the median global abundance of all bird species (estimated from (*19*)). Trends represent median and 95% Credible Interval estimates from a hierarchical logistic regression model (averaged across trends from all species groups; **see Fig. 3**). Note opposing directionality of trends between relatively natural vegetation types (*purple*) and anthromes and recently disturbed ecosystems (*pink*).

Over a third (36.8%; 95%-CRI: 34.8% - 39.0%) of all bird species considered are more likely to be recorded per sampling site in anthromes than non-anthromes, with this percentage increasing with the global abundance of species (***Table 1***). For example, 51% (95%-CRI: 48.03%-53.61%) of the most abundant quartile are more likely to occur in anthromes, compared with 17.4% (95%-CRI: 12.5%-22.28%) in the least abundant quartile, representing an approximately three-fold higher chance of the most abundant species being more associated with human-modified environments. This rises to 57.3% (95% CRI: 53.5% - 61.1%) when considering bird species in the top 10% for global abundance, whereas only 12.9% (95% CRI: 6.5% - 21.0%) of the species in the bottom 10% for global abundance are more likely to be recorded in anthromes (***Table 1*** and ***fig. S4***).

**Table 1.** Summary statistics underlying posterior estimates for the percentage of bird species more likely to occur in anthromes than non-anthromes. Summary statistics are presented for all bird species recorded in PREDICTS (top row), and for species recorded in PREDICTS that are represented in the bottom/top 10-50% of global abundances (by cumulative 5% increments) across *all* species with global abundance estimates from **(*19*)**. Percentages of species more likely to occur in anthromes were computed as derived quantities from the posterior distribution a hierarchical logistic regression model. For the purposes of computing these quantities, sites within plantation forests, croplands, pasture, urban environments and young secondary vegetation were classified as occurring in anthromes, and sites within primary vegetation, mature secondary vegetation and intermediate aged secondary vegetation were classified as occurring in non-anthromes.

| Percentile Cutoff<br>( Below/Above ) | Bottom or Top | Percentile value for species median global abundance<br>( Median global abundance estimate ) | Number of species in PREDICTS database | % more likely to occur in Anthromes<br>( Median (CI) ) |
| --- | --- | --- | --- | --- |
| 100% (B) † | All Species | 1,618,744,682 ‡ | 2,578* | 36.79 (34.79 - 38.98) |
| 10% (B) † | Bottom | 2,905 ‡ | 62 | 12.90 (06.45 - 20.97) |
| 15% (B) | Bottom | 11,155 ‡ | 103 | 14.56 (08.74 - 20.39) |
| 20% (B) | Bottom | 38,014 ‡ | 137 | 16.06 (10.95 - 21.9) |
| 25% (B) † | Bottom | 75,898 ‡ | 184 | 17.39 (12.5 - 22.28) |
| 30% (B) | Bottom | 120,929 ‡ | 247 | 18.22 (14.17 - 23.08) |
| 35% (B) | Bottom | 179,919 ‡ | 333 | 20.12 (16.51 - 24.32) |
| 40% (B) | Bottom | 250,261 ‡ | 436 | 21.33 (17.89 - 25.00) |
| 45% (B) | Bottom | 342,606 ‡ | 551 | 21.42 (18.33 - 24.86) |
| 50% (B) | Bottom | 450,955 ‡ | 685 | 22.48 (19.56 - 25.69) |
| 50% (A) | Top | 450,954 § | 1,893 | 41.94 (39.67 - 44.32) |
| 45% (A) | Top | 596,286 § | 1,743 | 43.26 (40.96 - 45.67) |
| 40% (A) | Top | 798,383 § | 1,573 | 44.56 (42.02 - 46.98) |
| 35% (A) | Top | 1,064,335 § | 1,413 | 46.14 (43.52 - 48.69) |
| 30% (A) | Top | 1,415,430 § | 1,242 | 48.15 (45.41 - 50.81) |
| 25% (A) † | Top | 1,915,783 § | 1,039 | 51.01 (48.03 - 53.61) |
| 20% (A) | Top | 2,753,114 § | 844 | 53.44 (50.47 - 56.28) |
| 15% (A) | Top | 4,291,374 § | 650 | 55.23 (52.00 - 58.46) |
| 10% (A) † | Top | 7,380,260 § | 447 | 57.27 (53.47 - 61.07) |

\* The total number of species used to compute the percentage of species more likely to occur in anthromes is the subset of the 3,146 species used in the main analysis, excluding species which: i) were only recorded in an original study that only surveyed secondary vegetation of indeterminate age (n = 25), and; ii) were only recorded in studies that surveyed *either* anthromes *or* non-anthromes but not both (n = 543). † Estimate presented in main text. ‡ Species with a median global abundance estimate *below* this value are included in this percentile bin. § Species with a median global abundance estimate *above* this value are included in this percentile bin.

Estimates of global abundances from different data sources (*19, 24*) are well correlated across species (r = 0.75; ***fig. S5***), which is unsurprising given that species’ global abundances vary over orders of magnitude (***Fig. 2***). As such, the result that relatively abundant species are more likely to be associated with anthromes and less abundant species more likely to avoid them is qualitatively and quantitatively consistent when alternate global population estimates are used in the analysis (***figs. S6-S9*** and ***Tables S4-S5***).

Similar conclusions hold for different bird ‘trophic niches’ (*22*): invertivores (n = 1637 species), omnivores (n = 532), frugivores (n = 376), granivores (n = 193), nectarivores (n = 138), terrestrial and aquatic herbivores (n = 15 and 23 respectively), species that eat other vertebrates (vertivores; n = 116), scavengers (n = 5) and aquatic predators (n = 111) (***Fig. 3*** and ***fig. S9*** for trophic niches which include at least 5% of the global avifauna; ***fig. S10*** and ***Table S5*** also include trophic niches with <5% of species). In particular, trends were consistently positive for the four anthromes and disturbed (young secondary) ecosystems and negative for primary, mature secondary and intermediate aged secondary vegetation (***Fig. 3 & figs. S9-S10***). The only exception was frugivores in primary vegetation, where the median slope estimate approximated zero.

**Fig. 3.**
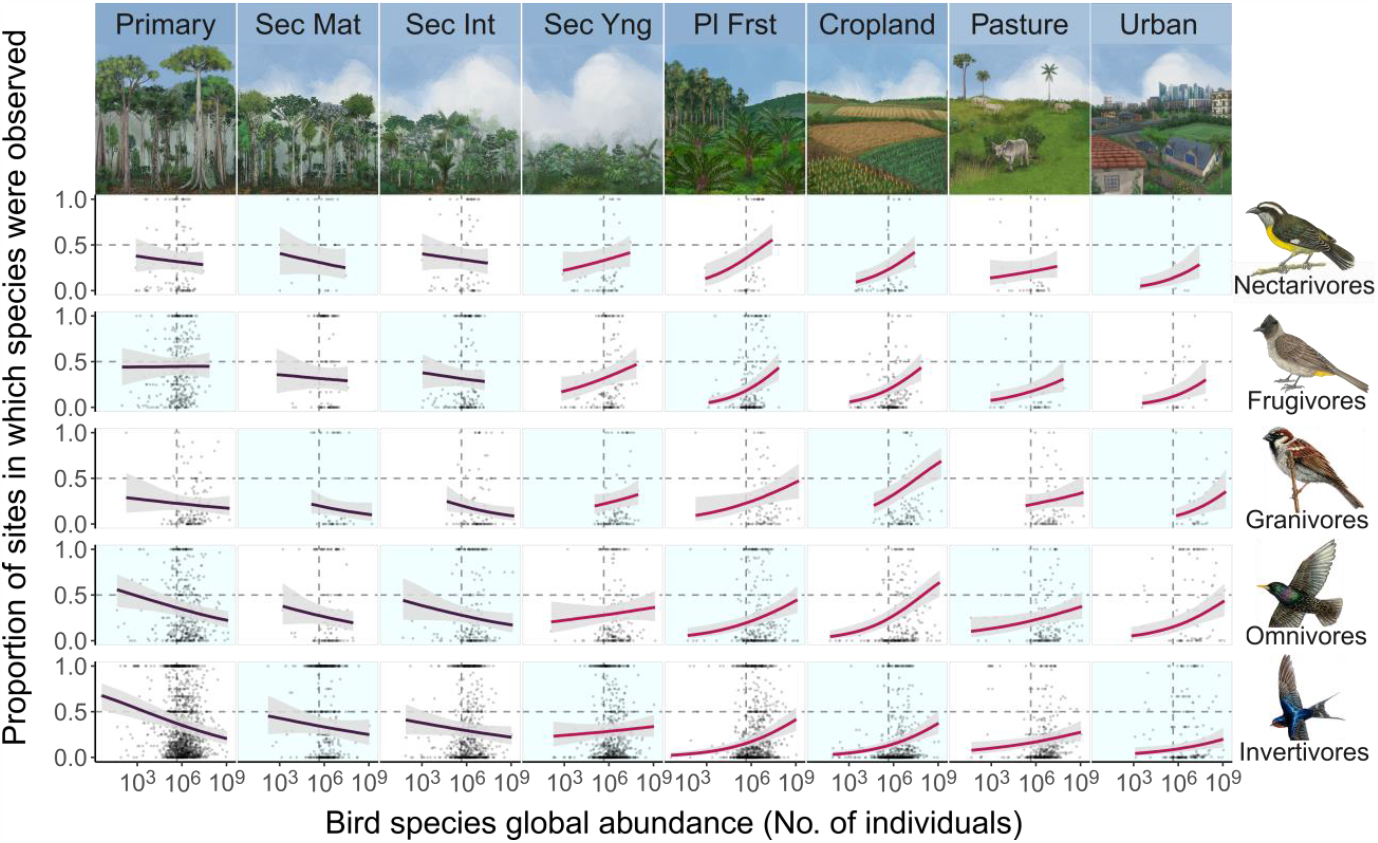
Trends in occurrence probability vs bird species global abundance for different trophic niches. X- and Y-axes, data points and dashed lines as in **Fig. 2**. Trends with 95% credible intervals were compiled from a hierarchical logistic regression model at the level of trophic niche within each land-cover type (*22*). Trends were fitted using species global abundance estimates from (*19*). Sec = secondary vegetation, Mat = Mature, Int = Intermediate aged, Yng = Young. Exemplar species (right) represent a globally abundant species from each trophic niche (Nectarivore (*Coereba flaveola*); Frugivore (*Pycnonotus barbatus*); Granivore (*Passer domesticus*); Omnivore (*Sturnus vulgaris*); Invertivore (*Hirundo rustica*)).

In addition to the overall association between species global abundance and anthrome use (***Figs. 2*** and ***3***), we also compared the likelihood of occurrence in primary vegetation to the likelihood of occurrence in the other land-cover types across the range of species global abundances (computed as contrasts between posterior occupancy trends in primary vegetation vs each other land-cover type; ***Fig. 4***). The results are consistent - on average, the more abundant a species is globally, the greater the likelihood that it will be observed in a given anthrome compared to primary vegetation, with little to no difference when comparing primary to mature and intermediate aged secondary vegetation.

**Fig. 4.**
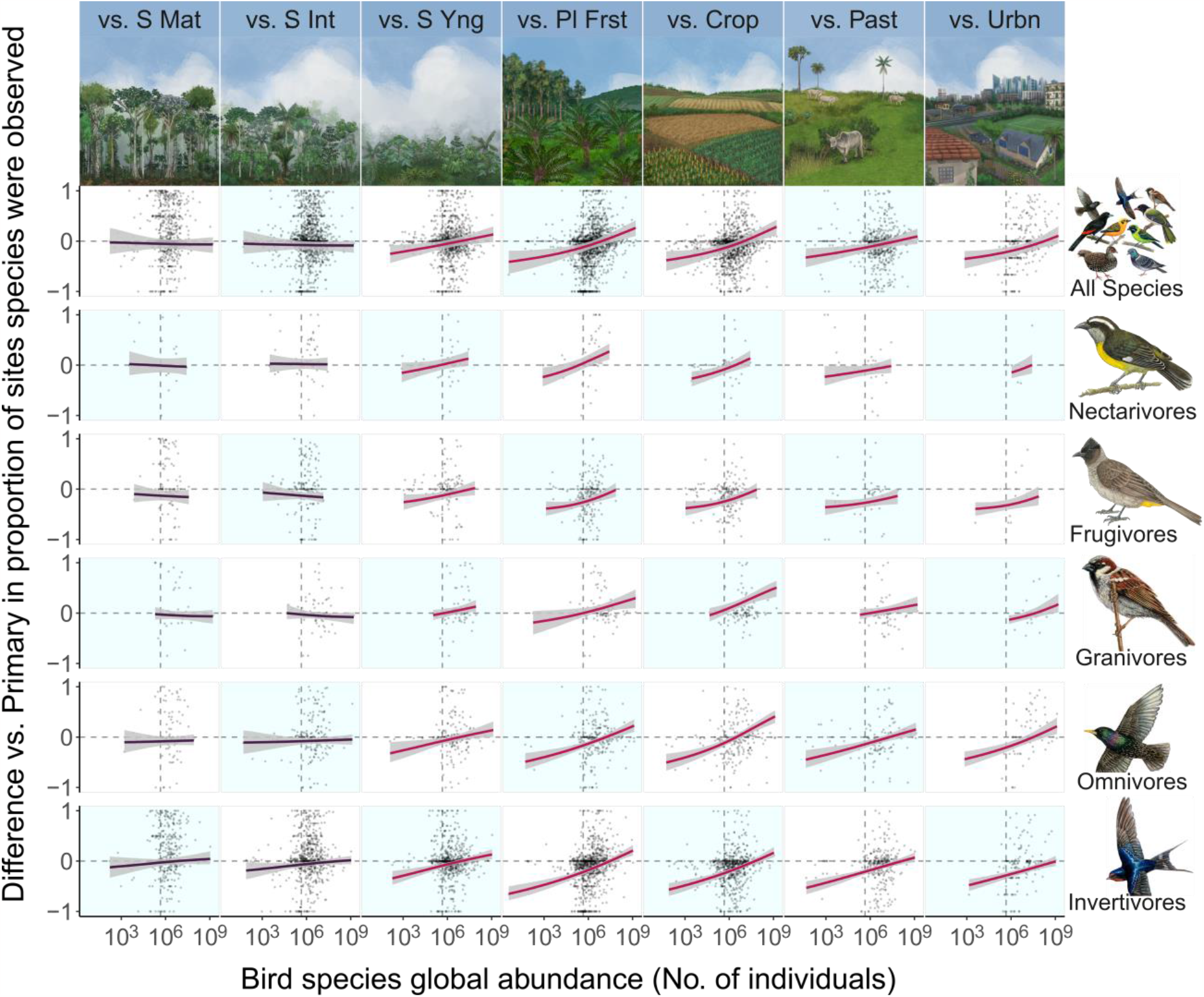
Likelihood of species occurrence in each land-cover type compared with occurrence in primary vegetation. Points represent empirical, species-specific differences in proportions of occupied sites (Y-axes), for each species recorded in at least one PREDICTS study that surveyed both primary vegetation and the corresponding land-cover type. S Mat = Secondary Mature vegetation, S Int = Secondary Intermediate, S Yng = Secondary Young. Dashed horizontal (0) lines indicate equal occurrence in primary vegetation and the comparison land-cover type; values <0 indicate higher occurrence in primary vegetation. X-axes denote species’ global abundance estimates (from (*19*)). Trends represent median and 95% credible intervals for contrasts computed from the posterior distributions of a hierarchical logistic regression model. Vertical dashed lines and exemplar species as in ***Fig. 3***.

The trophic groups do differ substantially, however, when we consider *absolute* differences in occurrence probability in primary vegetation compared with the other land-cover types. This can be seen via vertical ‘shifts’ of the trend lines relative to one another in ***Fig. 4***. In particular, granivorous species and the world’s more abundant omnivore species tend to have higher absolute frequencies of occurrence in young secondary vegetation and anthromes than in primary vegetation sites (i.e., the red lines are consistently above zero). As such, these two groups contribute disproportionately to the positive average contrast values for globally abundant species (top row of panels in ***Fig. 4***). This result adds context to a recent finding that plant and seed eating birds have consistently higher occurrence probabilities in anthromes than in primary vegetation, whereas results for omnivores were more variable (*25*), by showing that higher relative occupancy for omnivores is statistically dependent on species global abundance. By contrast, invertivore species were most often predicted to be less likely, on average, to occur in anthromes vs primary vegetation, in agreement with previous studies (*25*–*29*), across most of the range of global abundances. Frugivores were the only trophic niche for which species were predicted to occur most frequently in primary vegetation irrespective of the global abundances of species (***Fig. 4***; median trends less than zero for nearly all global abundance / vegetation combinations) – in line with previous studies indicating negative responses of frugivorous birds to human habitat modification (*25, 27, 28*). Despite differences in absolute levels of occurrence probability, the tendency for more globally abundant species to have increasing likelihoods of occurrence in anthromes relative to primary vegetation is consistent across feeding guilds (***Figs. 3-4***).

### Variation among species

While there are some differences among the different trophic niches (***Figs. 3-4***), there is much greater variation among the individual species within each niche type. Some individual species of all trophic niches are more likely, and others less likely, to occur in a given anthrome compared to ‘primary’ vegetation. Other factors such as size, morphology, additional niche specialisations and dispersal ability will also be important predictors of the sensitivity of species to habitat change (*27, 30*), as well as to species global abundances and population densities (*12, 19, 31*). For example, species that inhabit geographically extensive primary habitats, such as boreal forests may have high global abundances, regardless of their capacity to occupy human-modified ecosystems. Northern Eurasia, in particular, has high average bird range sizes (*31*) and extensive boreal forest (*32*) which has developed in the post-glacial period. Some of the variation among species will also be associated with sampling differences and accuracy, though this was taken into consideration within the statistical modelling by using site-, study- and species-level random effects. Raw data points summarising species proportions of occurrence shown in ***Figs. 2-4*** do not account for this complex structure but are shown here for transparency. Nonetheless, the overall pattern of a positive association between the global abundances of species and their capacity to inhabit anthromes emerges throughout the data.

An interesting question is where today’s globally abundant, human associated species came from, in the sense of what ecosystem types they previously inhabited. Some of today’s globally abundant species may, for example, have previously been restricted to marginal areas, ecotones or natural clearings and tree fall gaps in areas that were previously forested; in some instances human activities and livestock may have generated disturbances that resemble the previous activities of extinct megafauna. Results for young secondary vegetation are consistent with the contention that many human-associated species might previously have occupied disturbed environments, as young secondary trends are very similar to those for the four anthromes considered here (***Fig. 2-4***). Nonetheless, there is no universal set of adaptations or origins that predispose a species to occupy anthromes and become globally abundant – it is unlikely that any single origin would apply to all 37% of bird species estimated to be more likely to occur in anthromes than in more natural ecosystems. Some species may be more adaptable (‘generalists’ or individually flexible, such as intelligent corvids) and thus able to cope with human induced changes and novel environments (*33*), while others likely represent species that were in some sense ‘preadapted’ to the land-cover types human civilisation has created and spread (*11*). A level of pre-adaptation is suggested by the relative success of some trophic niches (especially granivores and omnivores; ***Fig. 4***) more than others, and some species have become particularly successful in some anthromes (e.g., *Streptopelia chinensis* and *Troglodytes aedon*). Genetic and behavioural changes can also contribute to increased abundances and the colonisation of anthromes (*34*). In many cases the differential success of species in anthromes and more natural habitat types may be mediated through species interactions (e.g., via the presence, absence or density of competitors, predators or prey), as well as through more direct environmental selection.

Some globally abundant species that occur across the world’s cities (*35*) and other highly modified land-cover types are labelled ‘synanthropic’. The reasons are again diverse. Species such as *Columba livia* and *Hirundo rustica* can use the built infrastructure in a similar way to the cliffs and rocky environments they use in more natural areas. For *Passer domesticus*, estimated to be the world’s most abundant bird (*19*), built and agricultural environments may replicate some elements of the steppes of their origin. Grassland species such as *Sturnus vulgaris* forage on grassland in urban parks and gardens and take advantage of human food provision. All of these species were observed in higher proportions of urban sites than sites in primary vegetation. Of course, not all anthromes are equivalent; the globally abundant *H. rustica*, for example, was marginally more likely to be observed in urban environments than in primary vegetation, but even more likely to be recorded in young secondary vegetation.

Regardless of the mechanisms, it is clear that species which can take advantage of human created environments can become highly globally abundant. This is especially the case for species such as *P. domesticus, S. vulgaris* and *C. livia*, for which humans have also provided a direct means of dispersal, releasing them across the globe (*36*). However, species have also dispersed under their own volition; *P. domesticus* spread with human agriculture before known direct introductions (*37*) and *Streptopelia decaocto* has rapidly expanded across Europe from the edges of its native range in the last century and a quarter (*38*). We surmise that a high proportion of human-associated species spread into some human-modified ecosystems – and many likely expanded their ranges prior to their histories being documented – as people successively transformed ecosystems during the Holocene.

The clear pattern of globally rarer, endemic or small ranged species being more likely to be restricted to relatively natural ecosystems has already been widely documented (e.g., (*15, 39*– *41*)). Our results are consistent with these previous studies, with globally threatened species such as *Ptilopachus nahani* and the threatened island endemic *Oriolus crassirostris* (*24*), for example, having considerably lower probability of occurrence in croplands compared with primary vegetation. In contrast, some low (global) abundance species were more frequently observed in anthromes, such as the tanagers *Ramphocelus flammigerus* and *Tangara arthus* in croplands.

Less than half of the globe’s ice free terrestrial land surface remains in semi-natural or what has been termed ‘fully wild’ condition, with the extent of croplands and pasture increasing rapidly over the last few centuries (*13*). Our results indicate that these changes have generated Anthropocene winners, globally abundant species occupying anthromes, and also losers, increasingly rare species associated with relatively undisturbed ecosystems. However, the nature and management of anthromes is dynamic, in line with the increasing speed of technological change over recent decades and centuries. Reported declines in widespread species that were once common (*42*–*44*), many of which were only common because they previously colonised anthromes that were managed according to more historical land-cover practices, indicate that these relationships may, in some instances, be transient. Changing human influences on global ecosystems mean that even some present-day ‘winners’ may become losers under future changing conditions (*45*), while some current ‘losers’ may also find new opportunities. Affinities, or lack thereof, of species with particular land use regimes are not necessarily static.

Whatever the particular evolutionary, ecological and human-association histories of each species, 57% of the world’s most abundant 10% of bird species are more frequent in anthromes than in more natural types of ecosystem. Given the environments that the majority of people live in, most of the birds that most of us see most of the time belong to these successful anthropophilic species. Maintaining the total abundance of the world’s birds and ecosystem functions associated with them requires consideration of human-modified environments, just as protecting the most localised species on Earth often focuses on the least-modified areas still remaining. This pattern is likely to generalise to many other taxa and provides both risks and opportunities for the management of biodiversity and ecosystem services.

## Acknowledgments

We would like to thank all the researchers who collected, calculated and curated the datasets used in this work. For the purpose of open access a Creative Commons Attribution (CC BY) license is applied to any Author Accepted Manuscript version arising from this submission. We thank Dr. Alexander Lovegrove for producing the scientific (ecosystem and species) illustrations which we have incorporated into ***Figs. 2-4***.

## Author contributions

Conceptualization: All authors

Methodology: All authors

Data curation: J.H.H., T.C. Formal analysis: T.C.

Writing - original draft, J.H.H., T.C.

Writing - Review & Editing:

All authors Visualization, T.C., J.H.H.

Funding acquisition, C.D.T.

## Materials and Methods

We used a global database of bird occurrences (*20*) and two different sets of published global bird abundance datasets (*19, 24*) to analyse species’ land-cover associations.

### Species Occurrence Data and Land Cover Associations

We obtained 395,305 detection/non-detection records for 3,146 bird species across 5,454 sample sites, representing more and less human-modified land use types from the PREDICTS database (*20*), from all 102 individual PREDICTS studies that surveyed birds. These studies surveyed an average of 2.25 land use types from 41 countries and were compiled from 88 original sources (*46*–*133*). We used the PREDICTS land-cover classification in our analyses, resulting in the categories of primary vegetation, mature secondary vegetation, intermediate aged secondary vegetation, young secondary vegetation, secondary vegetation of indeterminate age (*but see below*), cropland, pasture, plantation forest and urban. Primary vegetation constitutes area in which there is no record of historical human-induced habitat transformation, whereas secondary vegetation consists of sites in which the habitat has been transformed by humans in the past, but is not under contemporary human usage. For a full description of the PREDICTS land-cover type classifications, see (*20, 21*).

### Species taxonomy and trophic niches

Species names were matched between datasets using (*24, 134*–*137*). Synonyms were also identified using the R package taxize (*138*) based on itis.org (*139*). All species/country combinations present in PREDICTS (*20*) were checked (*24, 134*–*137*) and splits resolved based on location. Where the country in which the study was conducted was insufficient to resolve splits, the original citation underlying the PREDICTS data source (*see above*) was consulted to determine the location with species range maps (*140*). Coordinates were compared in QGIS (*141*) for the most complicated cases. Where species identity could not be resolved due to insufficient information, sympatric species splits in a given location or because the species is not known to occur in the recorded country (with no potential taxonomic revisions found) records were excluded.

We used a published classification to assign diet based trophic niches to each species (*22*). *Otus hartlaubi* was missing a classification so we categorised it as an invertivore based on closely related species. To compare the representativeness of bird species recorded in the PREDICTS database (*20*) with the wider global avifauna, we used published classifications of biogeographic realm (*22*) and habitat usage (*23*). Proportions of PREDICTS species are remarkably similar to analogous proportions for the entire global avifauna across biogeographic and functional categories (***Tables S1-S3***), suggesting that our analyses should allow us to make quite general inferences about the World’s birds.

### Global Bird Abundance Estimates

Global abundance estimates were obtained from (*19*) for 3,146 species for the main anlysis, and from (*24*) for 798 species (which also had estimates from (*19*)) to test the sensitivity of our results to different estimates of global abundance. Where only a range was provided in (*24*), rather than a mean estimate, the midpoint of the upper and lower values was used. Abundance estimates were highly correlated between the two data sources (r = 0.75; ***fig. S5***). Bird species represented in the PREDICTS (*20*) studies were broadly representative of the entire range of species global abundances (***figs. S1-2***), despite being unevenly distributed across the global abundance distribution, with globally rarer bird species less likely to be recorded.

### Statistical Model

We used a Bayesian hierarchical logistic regression model to assess whether the world’s most abundant bird species are predominantly associated with anthropogenic land-cover types. For our response variable, we converted the records extracted from PREDICTS (*20*) (many of which are abundance records) to detection/non-detection (1/0) values for each bird species within each sampling site (*Detect*\*NonDetect*_*i*_ in the model formula below). A record of 1 indicates that at least one individual of a particular species was detected at a given site during a PREDICTS study. A record of 0 indicates that no individuals of that species were detected at that site during the study, *but the species is known to occur in this region and is detectable using the same sampling methods*, as it was recorded in at least one site of any land-cover type during the same study.

To model the probability of detecting a bird species in a particular land-cover type as a function of species global abundance, we specified fixed effects to estimate land-cover specific intercept 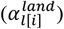 and slope 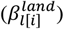 parameters with (scaled) log-global-abundance as a continuous explanatory variable, allowing species-level estimates to vary within and between land-cover types using random intercept offsets for species and species/land-cover interactions 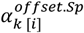 and 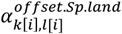 respectively). The random offset for the interaction between species identity and land-cover type 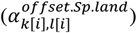 is important because the explanatory variable “global abundance” is a species-level variable (i.e, one value per species), for which we are interested in estimating land-cover-specific slope parameters 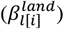. As such, the main effects of interest (effects of global abundance on occurrence within land-cover types) are slopes *across* species *within* land-cover types (i.e., estimated at the hierarchical level of “species within land-cover types” in our multilevel/hierarchical model), meaning that the ‘data’ to which the model fits these slopes are variable species-level intercepts estimated by the model (*142*). If we did not include the variable interaction term 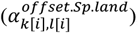, the model would estimate the 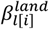 slopes under the implicit assumption that each species has the same occurrence probability within all nine land-cover types, thus leading to a mis-specified fit between model and data. Indeed, posterior predictive checks from an earlier iteration of the model excluding the species/land-cover interaction term demonstrated a much poorer fit and vastly exaggerated estimates of the 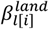 parameters (i.e., much more positive slopes in anthromes and much more negative slopes in non-anthromes, both of which were overconfident in terms of credible intervals, and ill-fitting).

We also specified (correlated) random intercept and slope offsets to account for variation among species assigned to different trophic niches (*22*) 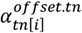 and 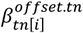, as well as (correlated) random interaction terms between trophic niche and land-cover type 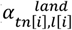 and 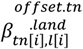. This allowed us to estimate separate trends across (scaled) log-global-abundances of species belonging to each trophic niche within each land-cover type, as well as estimating the aggregate land-cover level trends, which are effectively the weighted average across trophic niches within each land-cover type under this model specification (i.e., the fixed effect slopes 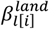. Covariance and correlation matrices are denoted as **S** and **R** respectively in the model formula below. Finally, we used random offsets to adjust for differences (e.g., sampling differences) across studies and sites 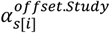 and 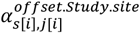.

The model was structured as follows:

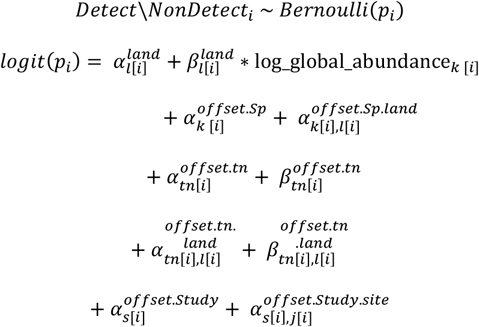

Subscripts in the model formula above refer to different hierarchical levels within the data and models. The number of categories at each of these hierarchical levels for the model fit using global abundance estimates from (*19*) are as follows: *i* = 1:395,305 rows of raw data; l = 1:9 land-cover types; *k* = 1:3,146 bird species, *tn* = 1:10 trophic niches, *s* = 1:102 PREDICTS studies, and *j* = 1: 5,454 sampling sites (corresponding numbers of hierarchical levels for the model fit using Bird-Life global abundance estimates (*24*) are presented in ***Table S6***).

We selected weakly regularising priors for each model parameter, using a ‘common sense’ approach (i.e., excluding impossible or wildly improbable parameter values), broadly following (*143*). The priors were as follows:

Fixed effects:

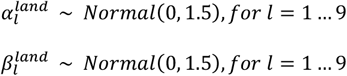

Random offsets and associated variance parameters:

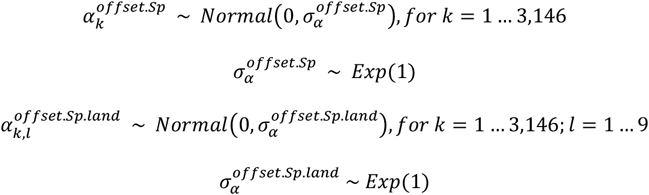

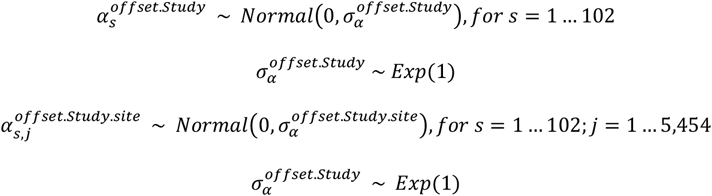

Random offsets with correlated intercepts and slopes:

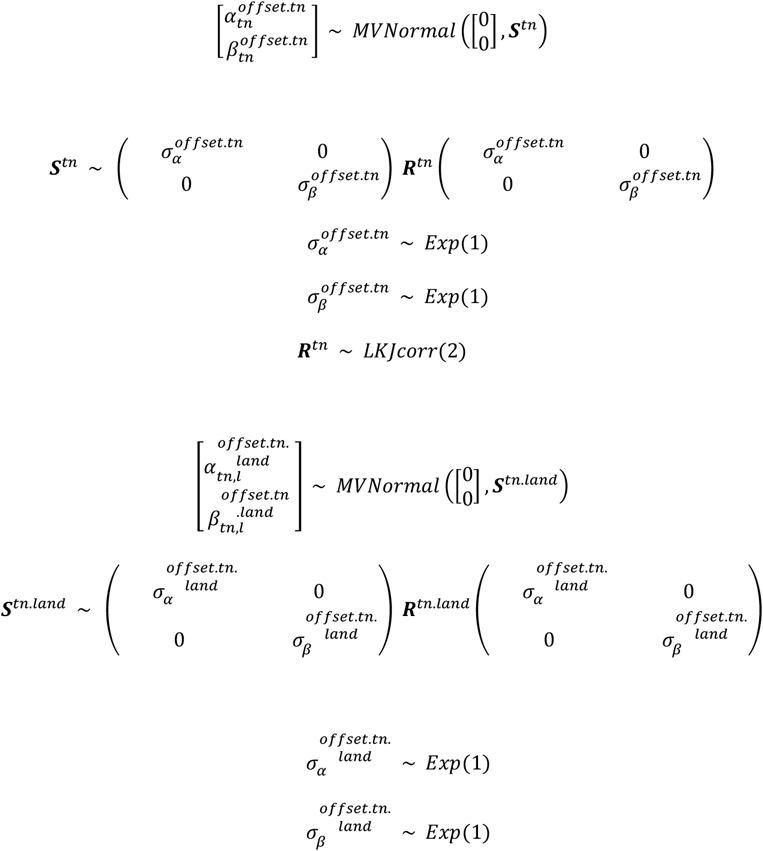

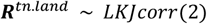

The model was fitted in R version 4.2.1 (*144*) using stan statistical programming language (*145*) via the R package brms (*146*). We used four MCMC chains of 24,000 iterations each, 12,000 of which were warmup. Chains were checked for convergence visually and using the Gelman-Rubin convergence statistic (*147*).

### Secondary vegetation of indeterminate age

We fit the hierarchical models to data including all PREDICTS land-cover types, including secondary vegetation of indeterminate age. However, results from this land-cover type are not directly useful in answering the overarching question underlying this paper, as the ordinal placement of secondary-indeterminate sites with respect to the recency of human land-cover transformation is not clear (i.e., sites may have been converted from primary vegetation in the recent past or a long time ago). However, we still included these data in the models because they are valuable in terms of estimating random offsets and variance parameters for species-, trophic-niche- and study-level random effects. Land-cover and trophic-nice/land-cover level trends for secondary vegetation of indeterminate age are presented in ***figs. S11-12***.

### Estimating the Percentage of Species that Occur More Frequently in Anthromes

Using the joint posterior distribution of the hierarchical logistic regression model, we derived the percentages of species estimated to occur more frequently in anthromes than non-anthromes for bird species in the PREDICTS database represented in the top and bottom 10-50% of global abundances across all species with abundance estimates from (*19*) by 5% increments (i.e., PREDICTS species in the top/bottom 10%, 15%… 50% of species global abundances across all 9,700 species from (*19*)), and for all species recorded in the PREDICTS database. This allowed us to make statements (with associated uncertainties) of the form of “*57% (95% CI:* 53.47% - 61.07%*) of bird species in top 10% of global abundances across <u>all</u> species are estimated to occur more frequently in anthromes than non-anthromes*”. Species were assigned to percentile bins for global abundance according to median global abundance estimates from (*19*). Of the 9,700 species with global abundance estimates, 685 species represented in PREDICTS that were sampled for in at least one anthrome and one non-anthrome site were in the bottom 50% and 1,893 were in the top 50% with respect to overall global abundances (see ***Table 1***. for sample sizes across the other percentile bins). For the purposes of this computation, sites within plantation forests, croplands, pasture, urban environments and young secondary vegetation were classified as occurring in anthromes. Sites within primary vegetation, mature secondary vegetation and intermediate aged secondary vegetation were classified as occurring in non-anthromes.

We computed the percentages as derived quantities implied by the posterior distribution by performing the following steps. We first compiled marginal posteriors for the occurrence probability of each species, within each land use type in which it was sampled for, as the inverse logit of the linear predictor:

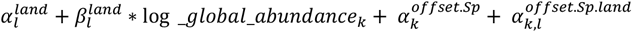

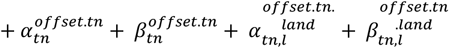

We then computed the mean occurrence probability for each species across the land-cover types for which it was surveyed in, within i) anthromes, and ii) non-anthromes, across each draw from the posterior distribution. Then, again at each draw from the posterior, we computed the percentage of species within each 5% top and bottom incremental band with a higher estimated probability of occurrence in anthromes vs non-anthromes, across all species represented in PREDICTS that were recorded in at least one PREDICTS study which surveyed an anthrome land use type and at least one study that surveyed at least one non-anthrome land use type. This gives us estimates of the percentages with associated uncertainties, which account for biases such as site- and study-level differences in sampling methods (as these are offset by site and study-level random effects). We performed bespoke posterior predictive checks to ensure that model estimates of these derived quantities did not deviate from the raw data (*see below*).

### Contrasts With Primary Vegetation

To assess whether, on average, species were more or less likely to occur in a given land use type than in primary vegetation, we derived contrast trends, with associated uncertainties, across the range of global abundances of species that were surveyed for in both primary vegetation and each of the other land use types. Contrast trends were computed as derived quantities across each iteration of the posterior distribution as:

1. Land-cover level contrasts representing differences in average occurrence probabilities between primary vegetation and each of the other land-cover types, computed as the difference between the inverse logit of:

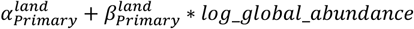

and

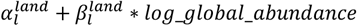

for each non-primary land use type, *l*, across the range of log global abundances for all species ‘surveyed for’ in both primary vegetation and land use type *l*.
2. Separate land-cover level contrast trends for each trophic-niche/land-cover combination, computed as the difference between the inverse logit of:

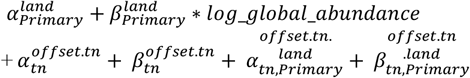

and

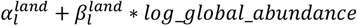

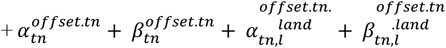

for each non-primary land use type, *l*, across the range of log global abundances for all species in a given trophic niche, *tn*, ‘surveyed for’ in both primary vegetation and land use type *l*.

In each instance, a value greater than zero indicates that species in this grouping with a global abundance value corresponding to a particular point along the contrast trend line are, on average, more likely to occur in the non-primary comparison land-cover type than in primary vegetation. A value less than zero indicates the converse.

### Model Validation

We performed posterior predictive checks to evaluate model fit. The motivating insight underlying posterior predictive checking is that one should be able to use parameter estimates from a model that provides an adequate fit to empirical data, in conjunction with the explanatory variables used to fit the model, to generate simulated datasets resembling the response variable (*148*–*150*). These posterior predictive simulations can be visually compared to various aspects of the empirical response variable to check for discrepancies that may indicate violations of model assumptions. Such comparisons can be made against the entire distribution of the response variable, or against bespoke test statistics computed from both the raw data and posterior predictive simulations (*151*). There is no general rule for choosing test statistics, as the most appropriate tests will depend on the particular analytical context. As recommended by Gelman *et al*. (*151*), we adopted a philosophy of “severe testing” with respect to model validation (i.e., designing tests which are highly likely to fail if a mis-specified model fit would result in misleading answers to important study conclusions (*152*–*154*)).

We performed a set of quite general tests to ensure that our model adequately reproduces the empirical response variable (detections/non-detections of birds) at various grouping levels that were and were not included in the model, as well as a set of more ‘bespoke’ tests to insure that the model can reproduce aspects of the data underlying inferences that are important to our specific conclusions, but which rely on quantities derived from the posterior distribution and are not explicitly included as explanatory variables. Together, these constitute a set of “severe tests” (*151*), ensuring both that our model provides a generally good fit to the data and that particular aspects of the data underlying specific conclusions are also well supported. We first produced 1,000 posterior predictive simulations for each row of data (n = 395,305) using the brms function “posterior_predict()” (*146*), and then computed test statistics using these simulations and the empirical response variable. We present plots of posterior predictive checks for the hierarchical logistic regression model fitted using species global abundance estimates from (*19*) (***figs. S13-23***), as this is the model upon which the main conclusions of our study are based.

For the general checks, we compiled test statistics constituting means (p) and standard deviations (√ p*1-p) of detection proportions, computed at grouping levels of land-cover type (***figs. S13-14***), trophic niches within land-cover types (***figs. S15-16***) and the countries in which detection/non-detections were recorded (***figs. S17-18***). Though ‘country’ was not included as a grouping variable in the model, including checks at this level allowed us to assess the possibility of geographic biases in our analysis (e.g., if the model provided a poor fit for countries with less data or in certain regions). Test statistics were computed over each of the 1,000 posterior predicted datasets and for the empirical response variable. At every grouping level, the model reliably reproduced the mean and standard deviation of the empirical response variable (i.e., test statistics computed from the empirical data sat perfectly within the distribution of test statistics computed over posterior predictive simulations; ***figs. S13-18***). Species-level test statistics, computed as the mean proportion of detections for each species within each land-cover type surveyed in studies which recorded that species, also reproduced the corresponding empirical proportions (***fig. S19***).

Given that the our aim is to investigate the relationship between the global abundance of species and their occurrence across different land-cover types, we also performed posterior predictive checks to assess whether there was any bias in model fit at particular points along the global species abundance distribution. This is important because we use the model to make inferences and comparisons with respect to species at the extreme ends of the global abundance distribution (e.g., species in the top and bottom 10% for global abundance), and poor fit for species with low or high global abundances could potentially invalidate such inferences. To do this, we subtracted raw empirical detection proportions for each species in each land-cover type (Y-axes; ***fig. S19***) from the same quantity computed from each posterior predictive dataset (Y-axes; ***fig. S19***), to compute the difference between the detection proportion the model implies for each species it’s observed empirical value. We then plotted these model departure distributions against species log-global-abundances (***fig. S20***). We applied the same procedure using empirical species/land-cover level contrast values and their counterparts computed over posterior predictive simulations (***fig. S21***). Both of these tests confirm that there is no bias in model performance at any particular point along the global abundance distribution, and in particular, that the model performs well with respect to inferences highly globally abundant and globally rare species.

Finally, we performed posterior predictive checks to ensure that derived quantities underlying the percentages of species more likely to occur in anthromes than non-anthromes are fully supported by the data. To do this, we computed the same percentages for the same bins of species within anthromes and non-anthromes (grouped by their position along the rank global abundance distribution) that we computed from model estimates (***Table 1***. and ***fig. S4***.), on raw empirical proportions of these quantities and proportions computed on posterior predictive simulations. Test statistics derived from the model’s posterior predictive distribution largely conform to the same quantity computed on the raw data (***figs. S22-23***). Indeed, the empirical quantities fall squarely within the distributions of test statistics for all PREDICTS species (***fig. S22***) and for all percentages computes for groupings in the bottom half of the global abundance distribution (***fig. S23***). However, for some groupings in the top half of the global abundance distribution (the top 15%, 20% and 25% bins), posterior predictive simulations slightly undershoot the empirical quantity, suggesting that the model may imply somewhat conservative estimates of these quantities (***Table 1***.). Overall, our hierarchical logistic regression model demonstrates a remarkably good fit to the empirical data.

**Fig. S1.**
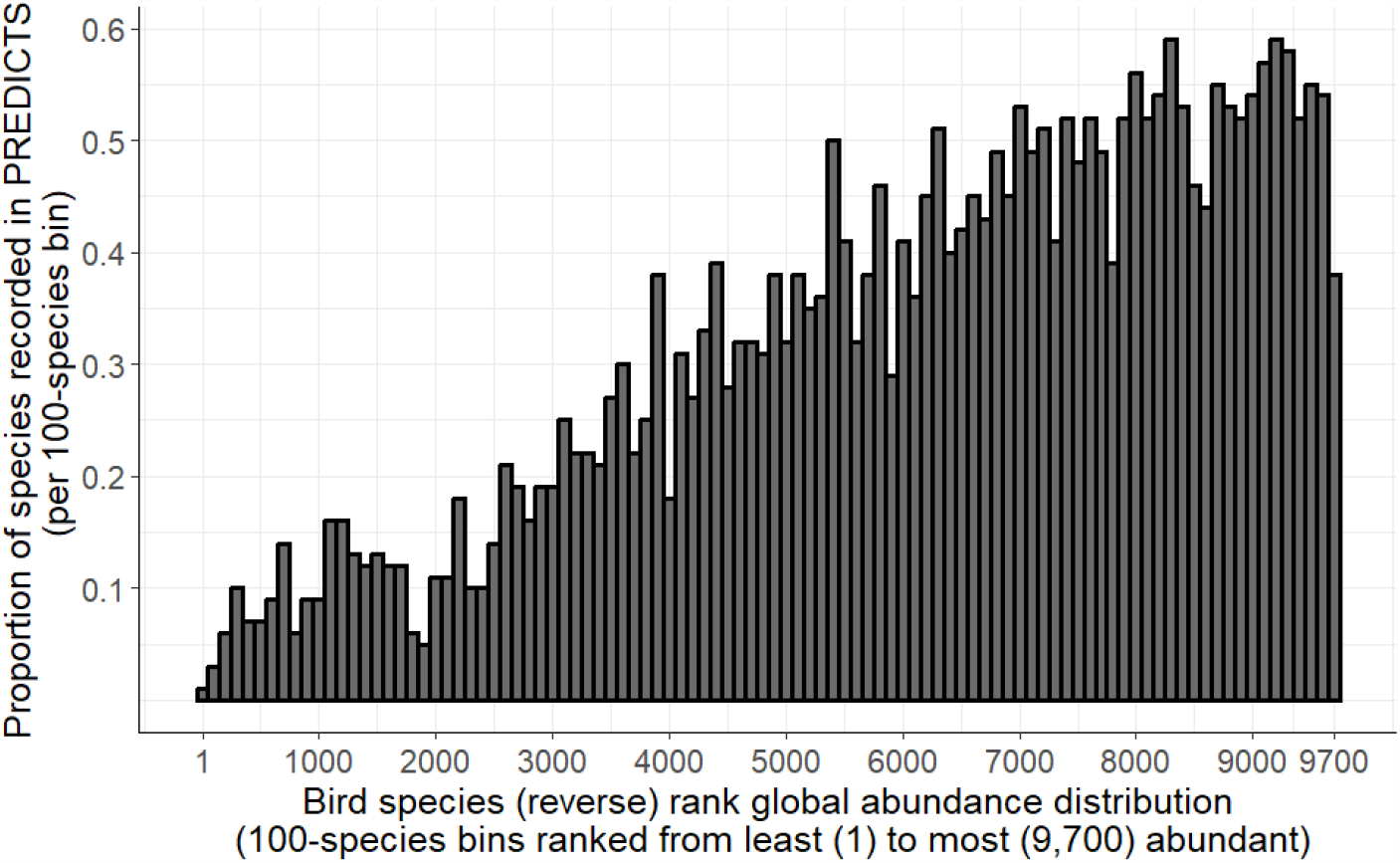
Proportion of bird species recorded in PREDICTS database (*20*) per 100 species along rank global abundance distribution. Each bin represents the proportion of species recorded in PREDICTS per 100 species ranked from lowest to highest median global abundance for all 9,700 species with global abundance estimates from Callaghan *et al*. (*19*). Globally abundant bird species are more likely to be recorded in PREDICTS, but species are well represented across the entire rank distribution of global abundances.

**Fig. S2.**
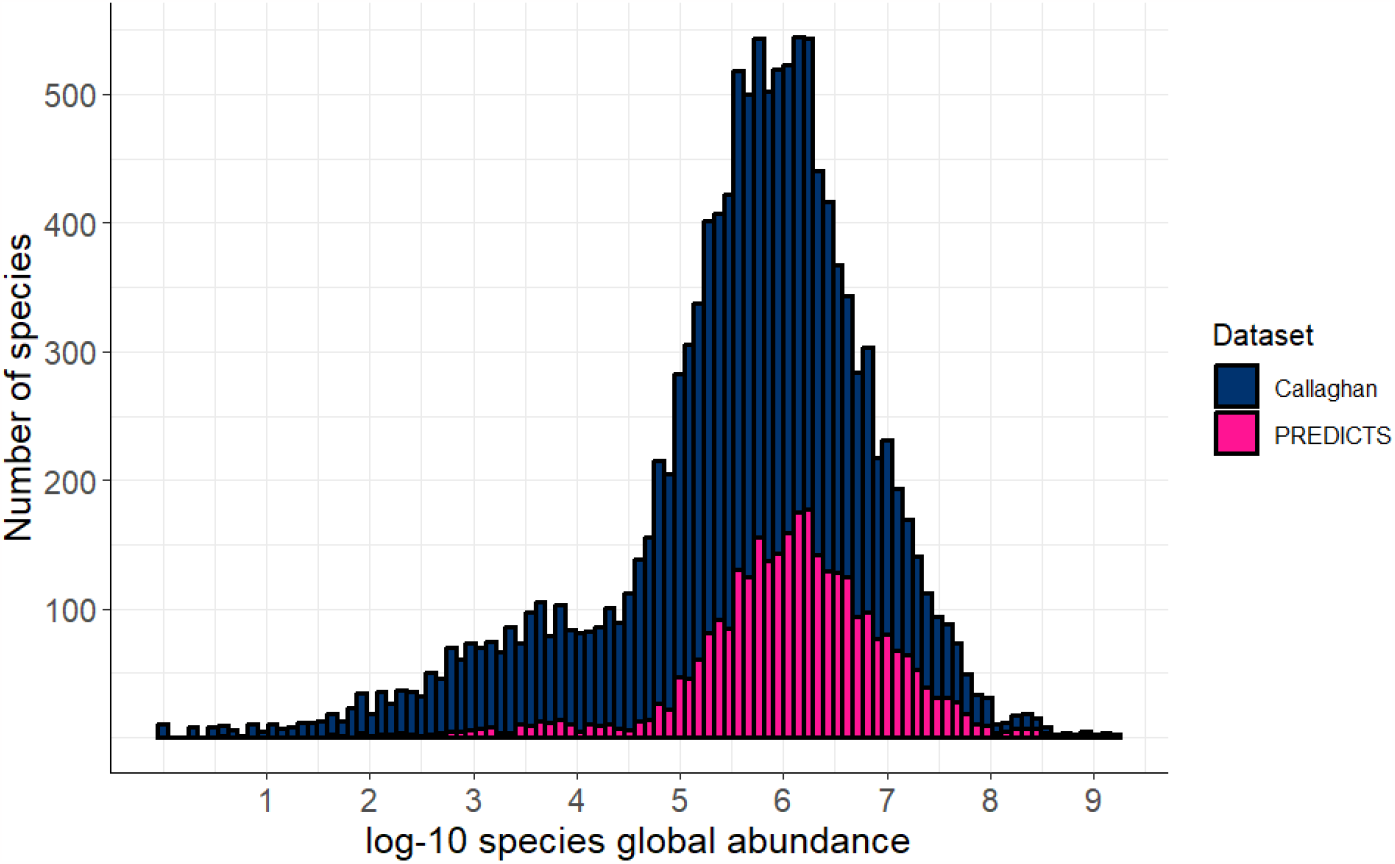
Number of bird species represented in Callaghan *et al*. (*19*) **and the PREDICTS database** (*20*) **vs. log-10 species global abundance**.

**Fig. S3.**
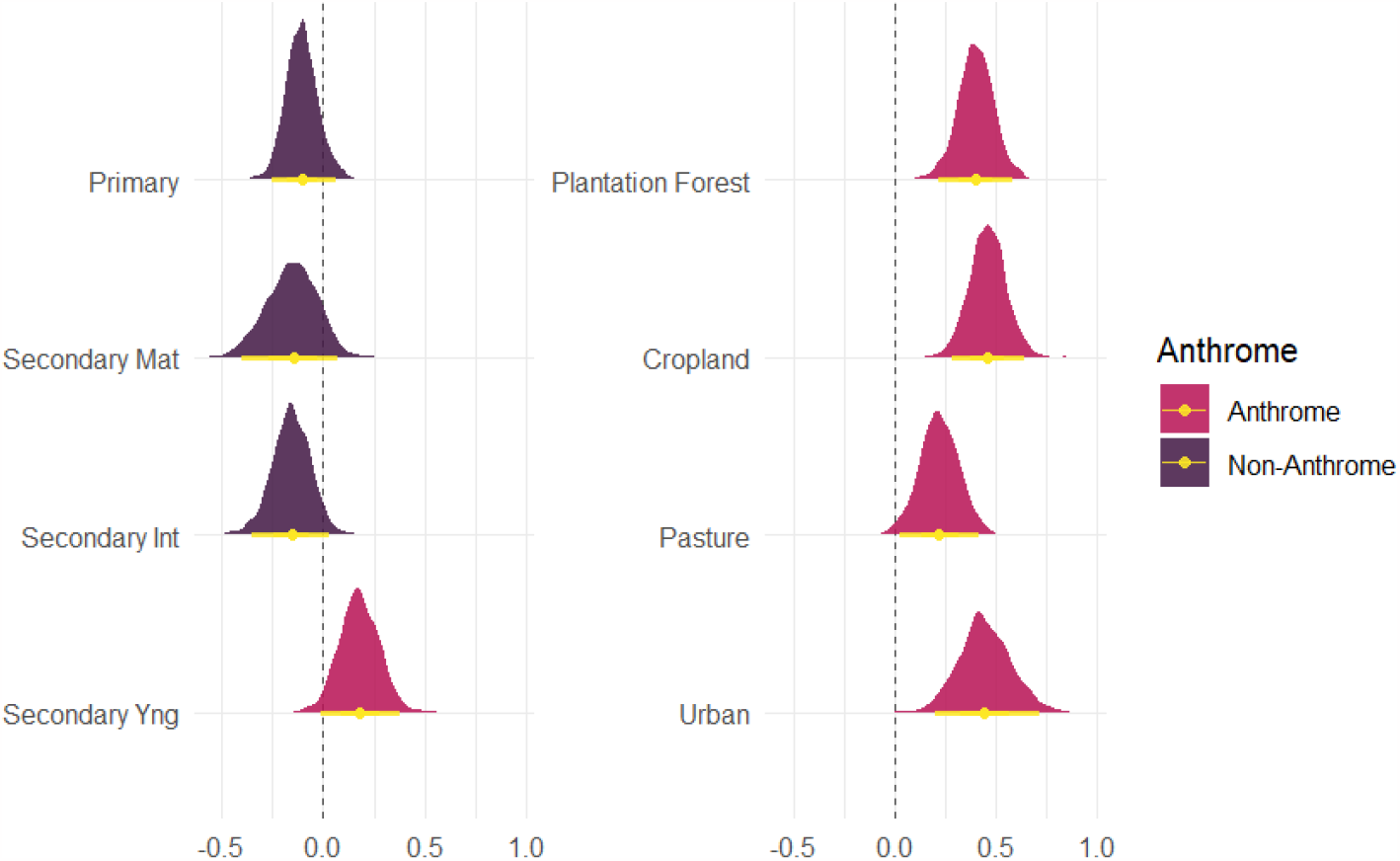
Posterior distributions for land-cover level ‘fixed effect’ slopes of (scaled) log global abundance on species occurrence from Bayesian hierarchical logistic regression model. Vertical dashed lines at zero represent ‘no effect’ of species global abundance on site-level occurrence within a particular land-cover type. Point intervals represent 95% Bayesian credible intervals. Note that slope estimates for non-anthrome land-cover types are lower in magnitude and, as a result, more uncertain in terms of directionality of effect compared with positive slopes in anthrome land-cover types. Parameter estimates are from model fitted using global abundance estimates from (*19*).

**Fig. S4.**
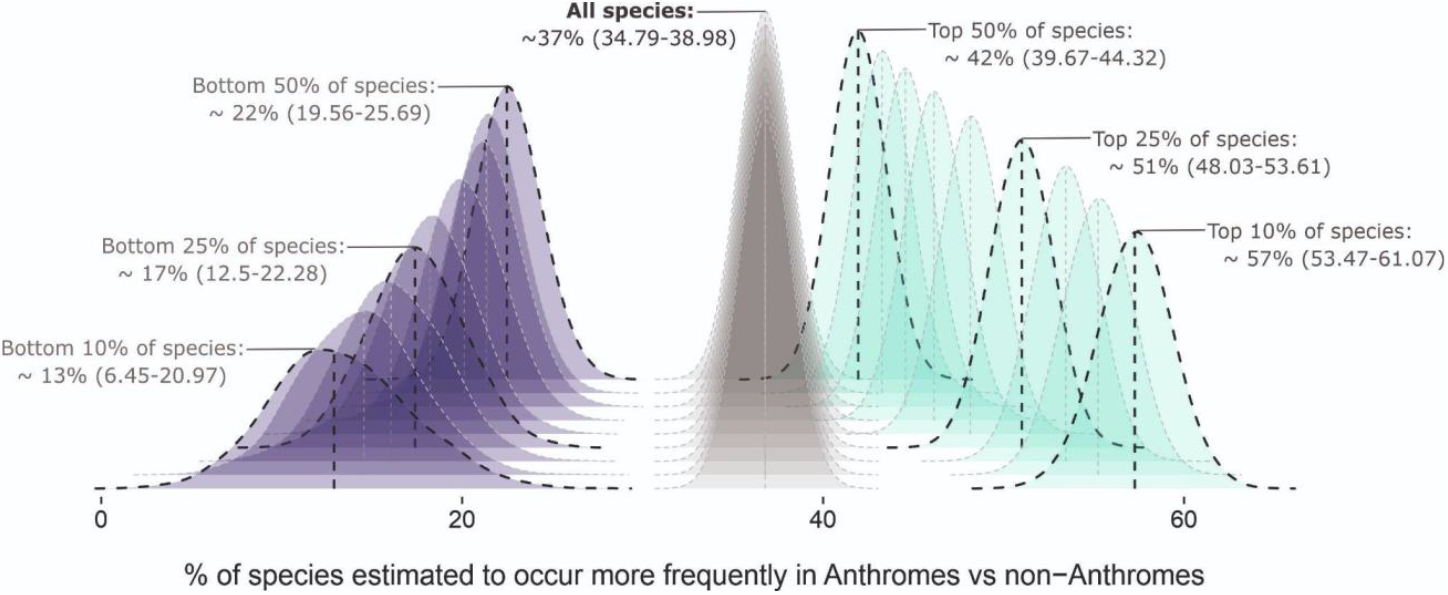
Posterior density ridge plot for the percentage of the species estimated to occur more frequently in anthromes than non-anthromes. Densities are shown for bird species represented in the PREDICTS database subset by global abundance estimates across *all* bird species with global abundance estimates from (*19*), for: **i)** Species that fall within the bottom 10-50% for global abundance (*left/purple*; by 5% increments); **ii)** All bird species in the PREDICTS database (*middle/grey*; replicated across each ridge), and; **iii)** Species that fall within the top 10-50% for global abundance (*right-turquoise*; by 5% increments). Vertical dashed lines denote the median value for each distribution. Median values and 95% Bayesian credible intervals (parentheses) are displayed as annotations for a subset of bottom/top density bins and for all PREDICTS species (*see* ***Table 1***. *for summary statistics for all density bins*). For the purposes of these computations, sites within plantation forests, croplands, pasture, urban environments and young secondary vegetation were classified as occurring in anthromes, and sites within primary vegetation, mature secondary vegetation and intermediate aged secondary vegetation were classified as occurring in non-anthromes.

**Fig. S5.**
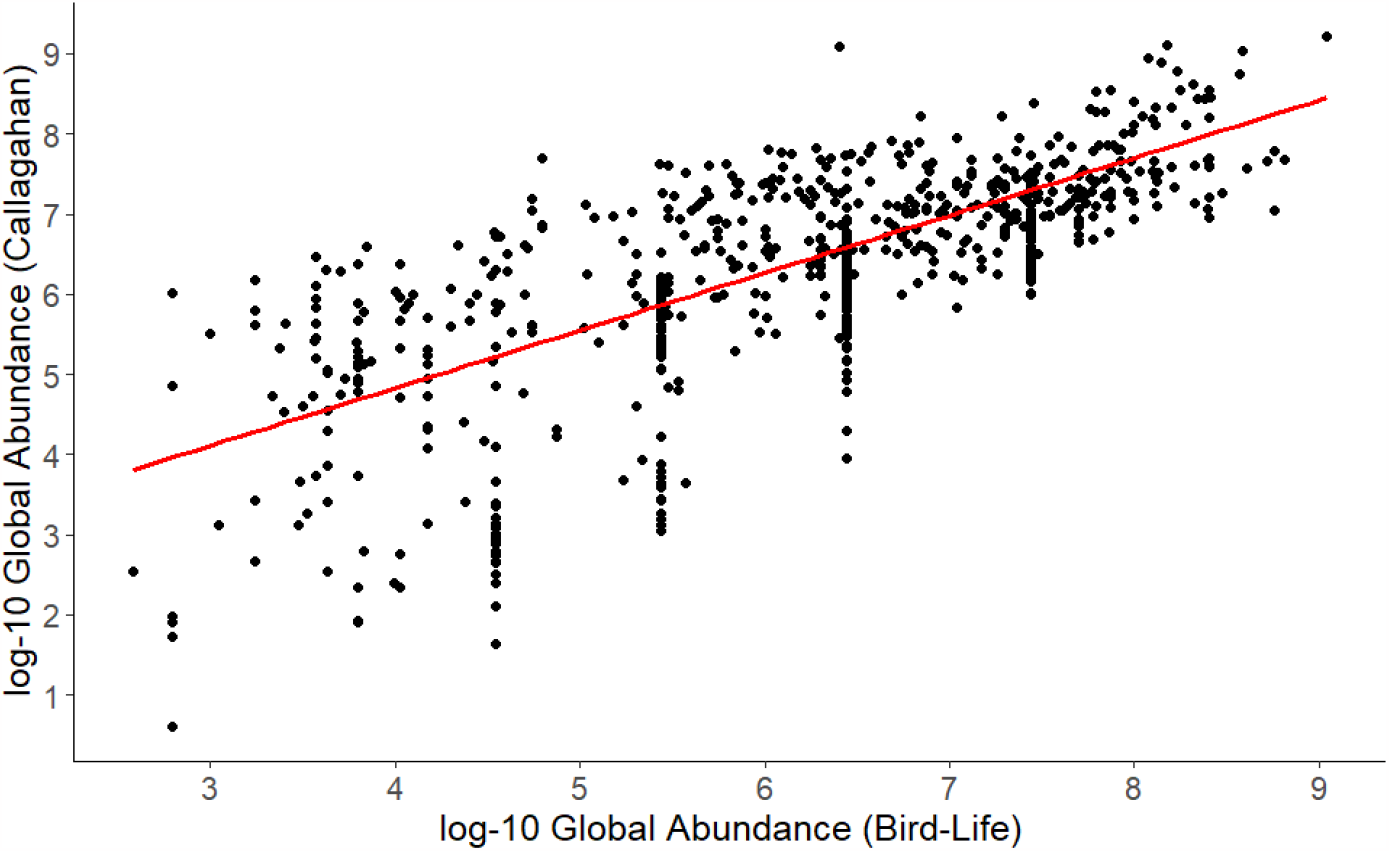
Scatterplot of log-10 median global abundance estimates sourced from Callaghan *et al*. (*19*) (y-axis) and BirdLife International (*24*) (x-axis) for all bird species in PREDICTS database with abundance estimates from both sources. Abundance estimates are well correlated across the range of global abundances (Pearson correlation coefficient of 0.75). Clustering in BirdLife estimates is due to the fact that many species are assigned the same estimated abundance range.

**Fig. S6.**
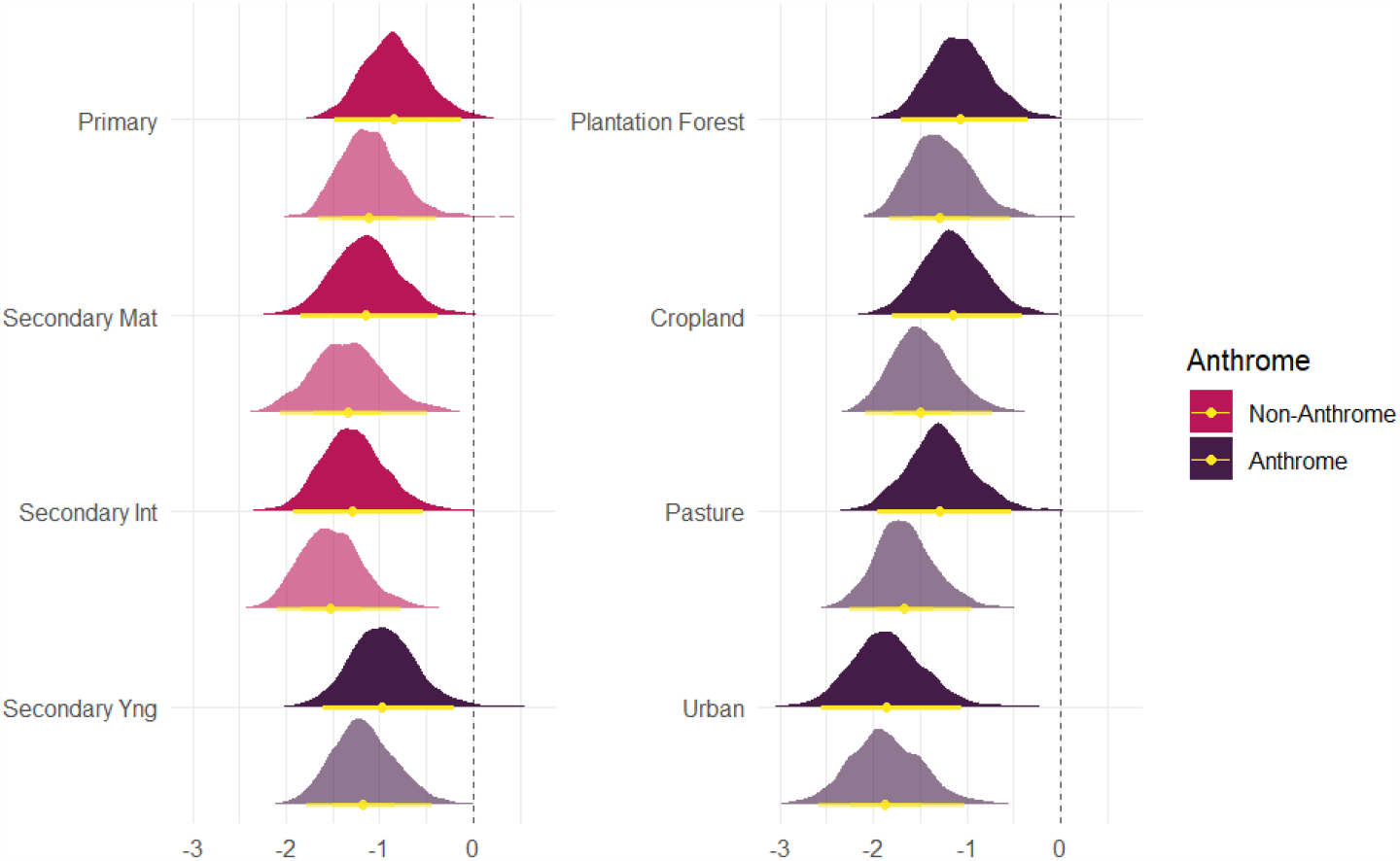
Posterior distributions for land-cover level ‘fixed effect’ intercepts (logistic scale) from Bayesian hierarchical regression models. Dark shaded distributions are parameter estimates from model using global abundance estimates sourced from Callaghan *et al*.(*19*), light shaded distributions are parameter estimates from model using global abundance estimates sources from BirdLife International (*24*). Vertical dashed lines at zero represent a 0.5 occurrence probability in a given land-cover type for species with mean global abundance. Point intervals represent 95% Bayesian credible intervals.

**Fig. S7.**
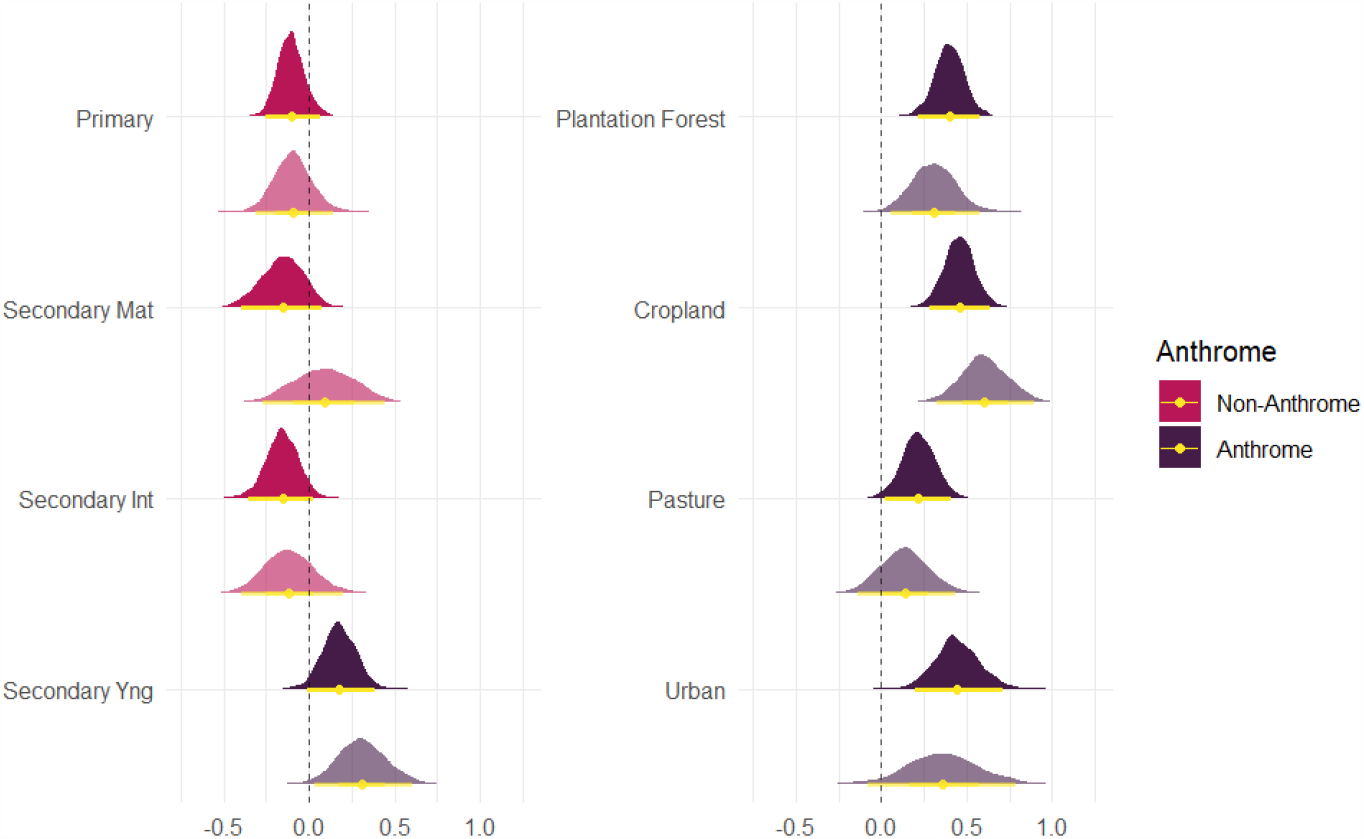
Posterior distributions for land-cover level ‘fixed effect’ slopes of (scaled) log global abundance on species occurrence from Bayesian hierarchical regression models. Dark shaded distributions are parameter estimates from model using global abundance estimates sourced from Callaghan *et al*.(*19*), light shaded distributions are parameter estimates from model using global abundance estimates sources from BirdLife International (*24*). Vertical dashed lines at zero represent ‘no effect’ of species global abundance on site-level occurrence within a particular land-cover type. Point intervals represent 95% Bayesian credible intervals.

**Fig. S8.**
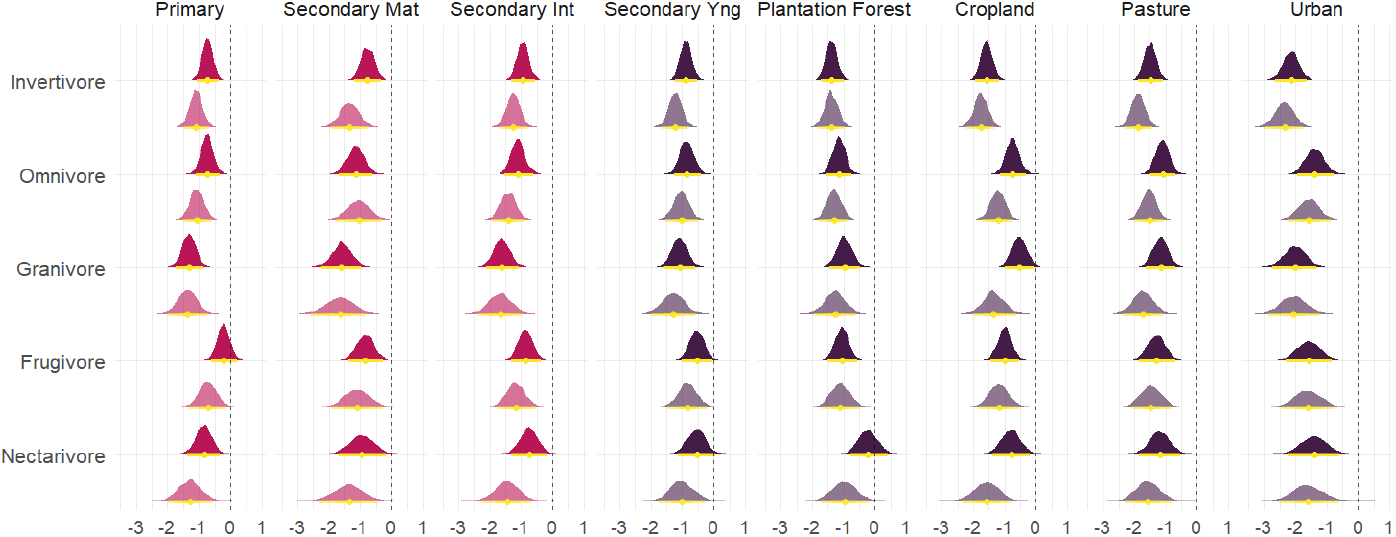
Posterior distributions for land-cover level intercepts for each trophic niche which includes at least 5% of the global avifauna (logistic scale) from Bayesian hierarchical regression models. Dark shaded distributions are parameter estimates from model using global abundance estimates sourced from Callaghan *et al*.(*19*), light shaded distributions are parameter estimates from model using global abundance estimates sources from BirdLife International (*24*). Vertical dashed lines at zero represent a 0.5 occurrence probability in a given land-cover type for species with median global abundance. Point intervals represent 95% Bayesian credible intervals.

**Fig. S9.**
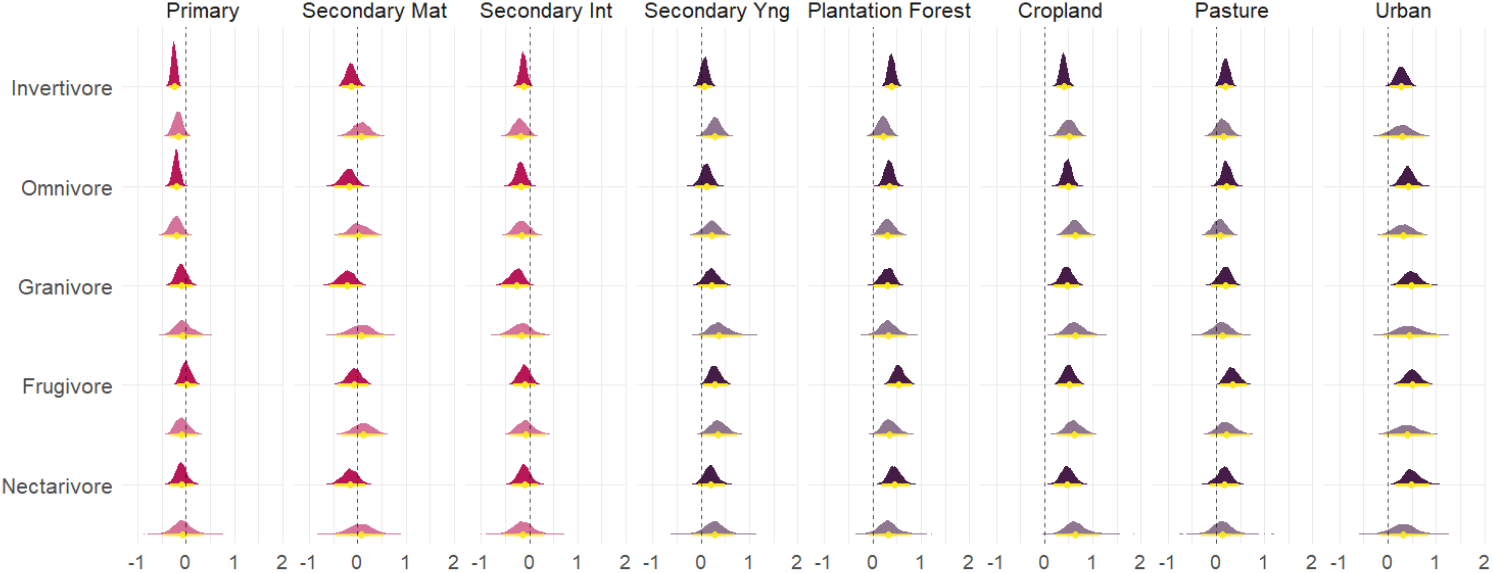
Posterior distributions for land-cover level slopes of (scaled) log global abundance on species occurrence for each trophic niche which includes at least 5% of the global avifauna from Bayesian hierarchical regression models. Dark shaded distributions are parameter estimates from model using global abundance estimates sourced from Callaghan *et al*.(*19*), light shaded distributions are parameter estimates from model using global abundance estimates sources from BirdLife International (*24*). Vertical dashed lines at zero represent ‘no effect’ of species global abundance on site-level occurrence within a particular land-cover type. Point intervals represent 95% Bayesian credible intervals.

**Fig. S10.**
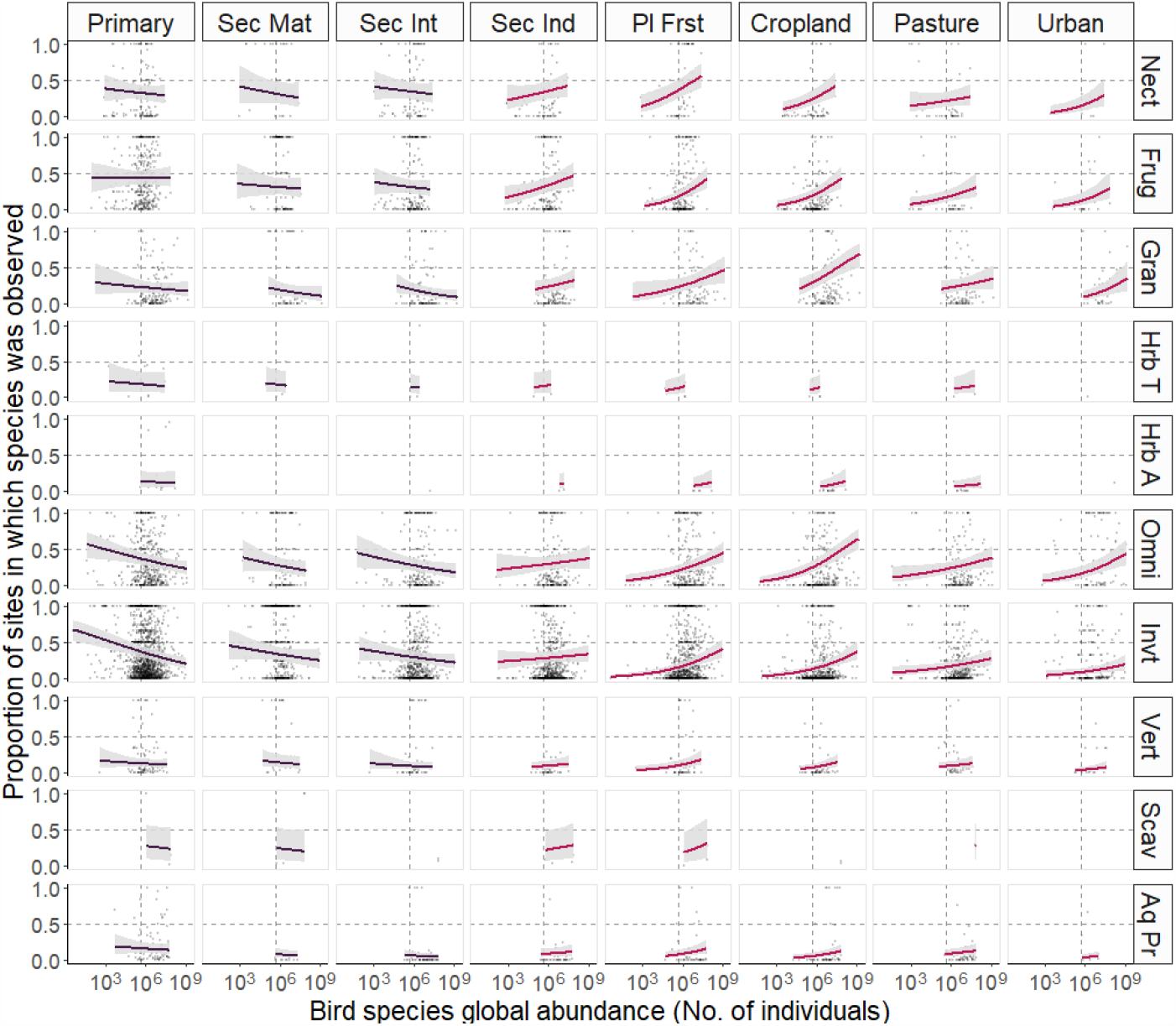
Trends in occurrence probability vs bird species global abundance for all trophic-niche/land-cover combinations. X-axes represent species’ global abundance estimates (*19*) and Y-axes show proportions of land-cover specific survey sites in which bird species were recorded. Points in each panel show empirical, species-specific proportions of sites occupied in a given land-cover type, for each species recorded in at least one PREDICTS (*20*) study surveying which recorded that species. Abbreviations: Primary (primary vegetation), Sec Mat (seconadry mature vegetation), Sec Int (secondary intermediate vegetation), Sec Ind (secondary vegetation of indeterminate age), Pl First (plantation forest), Nect (nectivore), Frug (frugivore), Gran (granivore), Hrb T (terrestrial herbivore), Hrb A (aquatic herbivore), Omni (omnivore), Invt (invertivore), Vert (vertivore), Scav (scavenger), Aq Pr (aquatic predator).

**Fig. S11.**
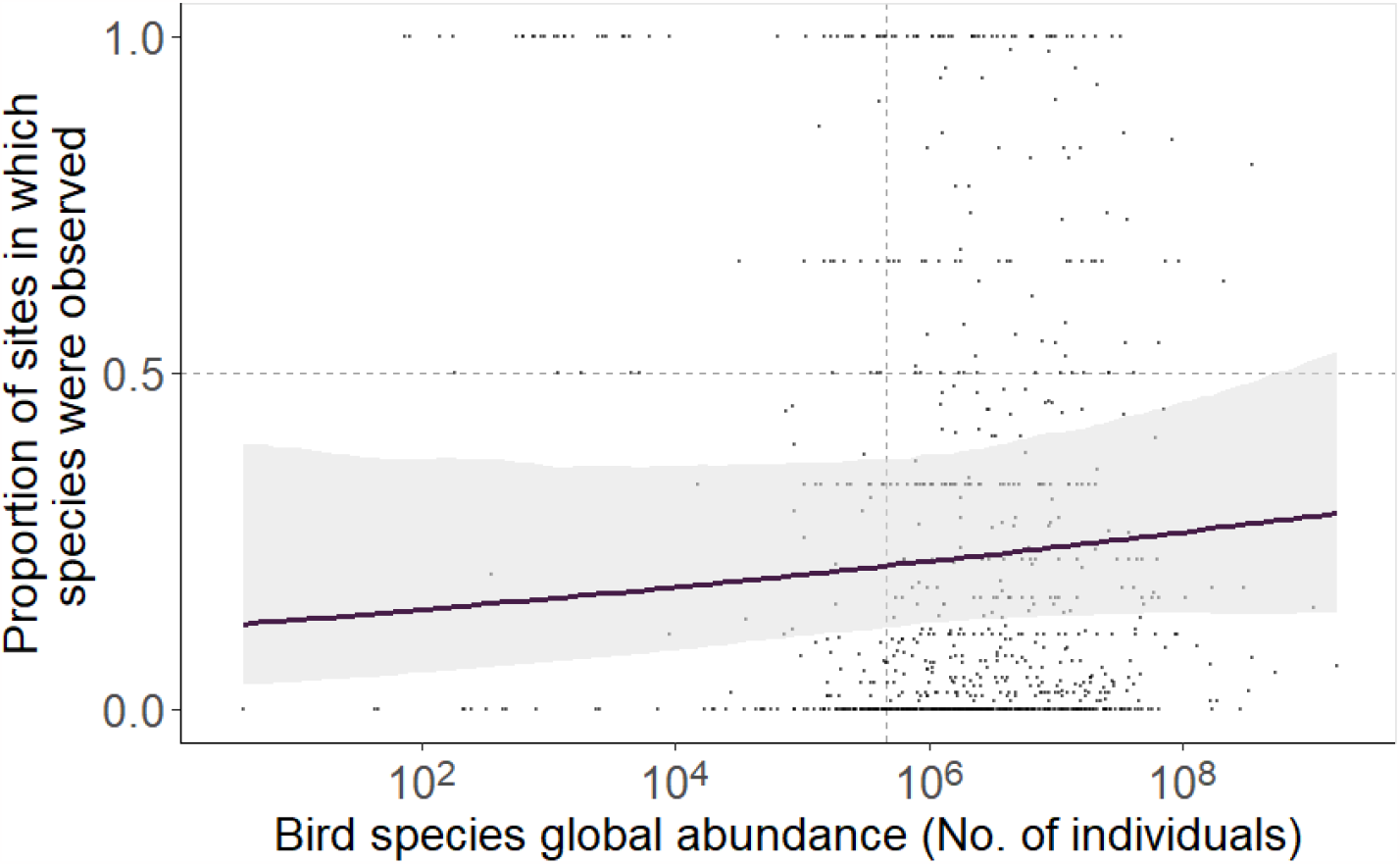
Land-cover level trend in occurrence probability vs bird species global abundance for secondary vegetation of indeterminate age. Points depict raw proportions of sites in secondary vegetation of intermediate age in which each species was recorded. Trend depicts the median and 95% Bayesian credible interval estimated from the hierarchical logistic regression model using global abundance estimates from (*19*).

**Fig. S12.**
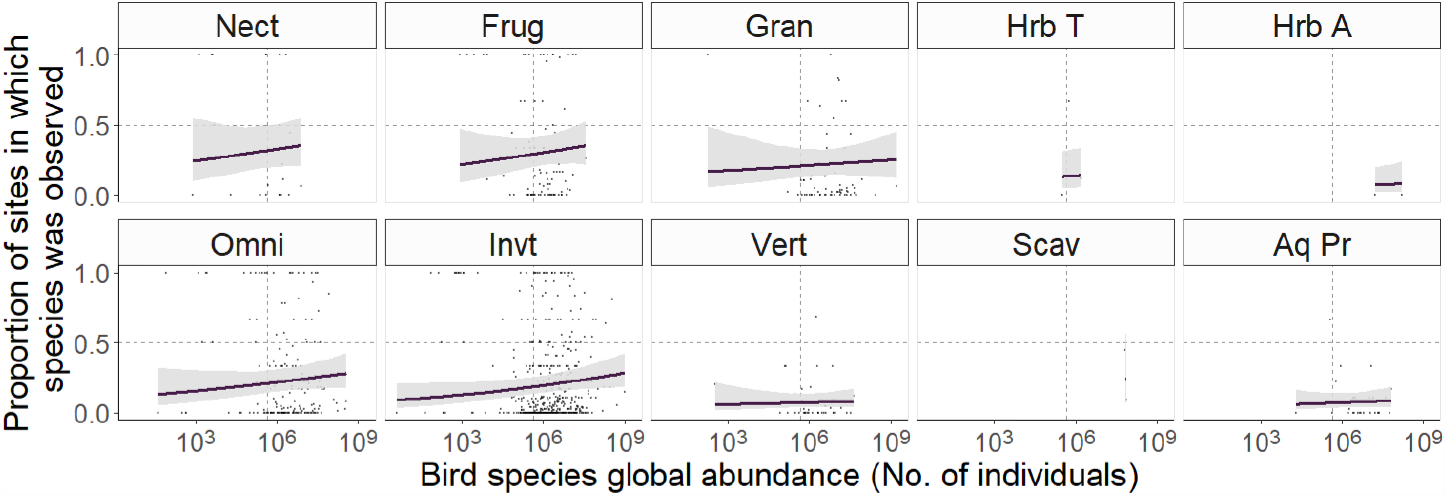
Trends in occurrence probability vs bird species global abundance for all trophic-niches in sites of secondary vegetation of indeterminate age. Points depict raw proportions of sites in secondary vegetation of intermediate age in which each species was recorded. Trends depict median and 95% Bayesian credible intervals estimated from the hierarchical logistic regression model using global abundance estimates from (*19*). Panel titles depict abbreviated names for trophic niches (Nect = Nectarivores, Frug = Frugivores, Gran = Granivores, Hrb T = Terrestrial herbivores, Hrb A = Aquatic herbivores, Omni = Omnivores, Invt = Invertivores, Vert = Vertivores, Scav = Scavengers, Aq Pr = Aquatic predators).

**Fig. S13.**
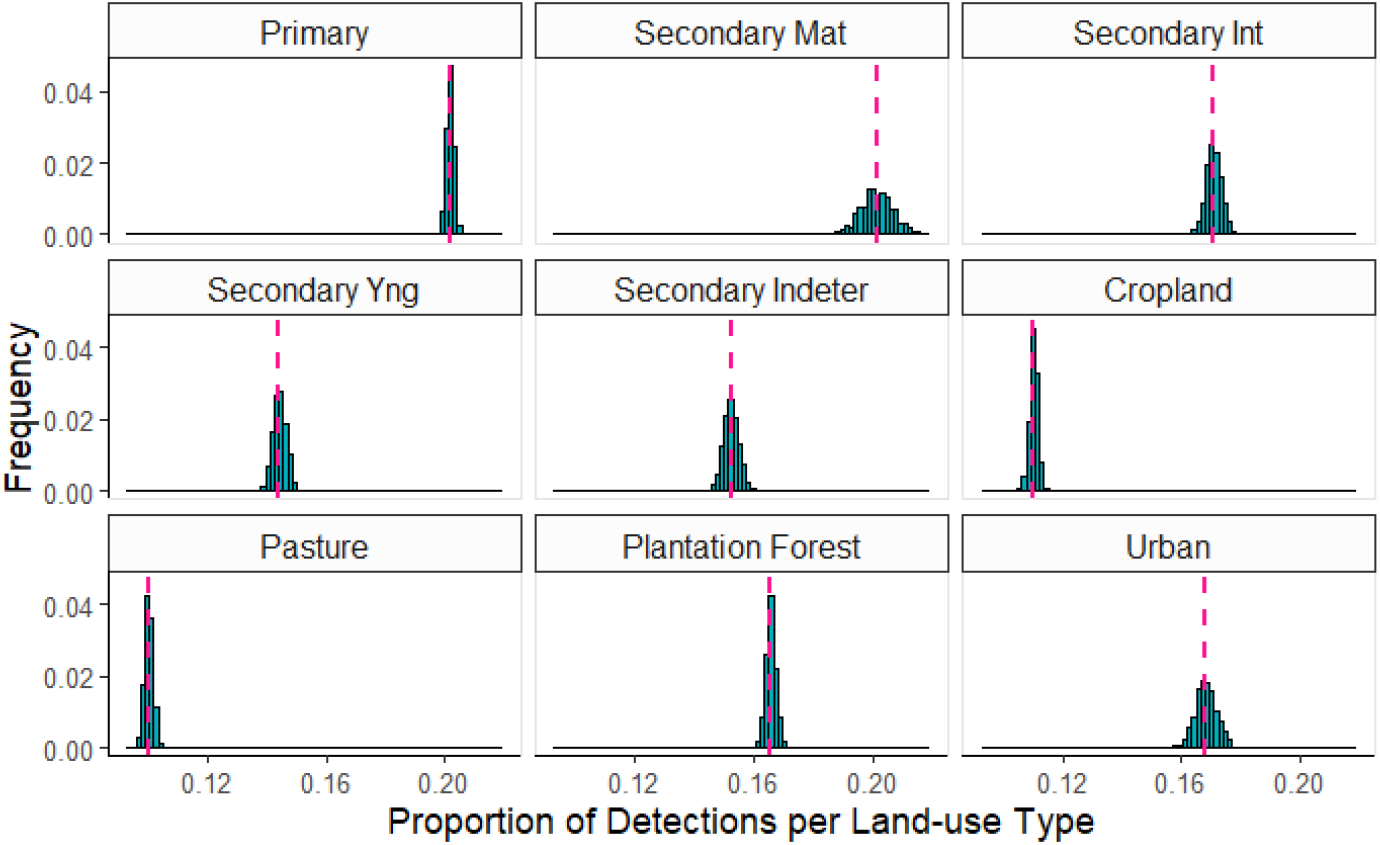
Posterior predictive check for proportion of species detections observed in each land-cover type. Histograms depict the distribution of test statistics (mean/proportion of detections) computed for each of 1,000 datasets simulated from the posterior predictive distribution of the Bayesian hierarchical logistic regression model. Vertical dashed lines depict the same test statistic computed from the raw empirical data.

**Fig. S14.**
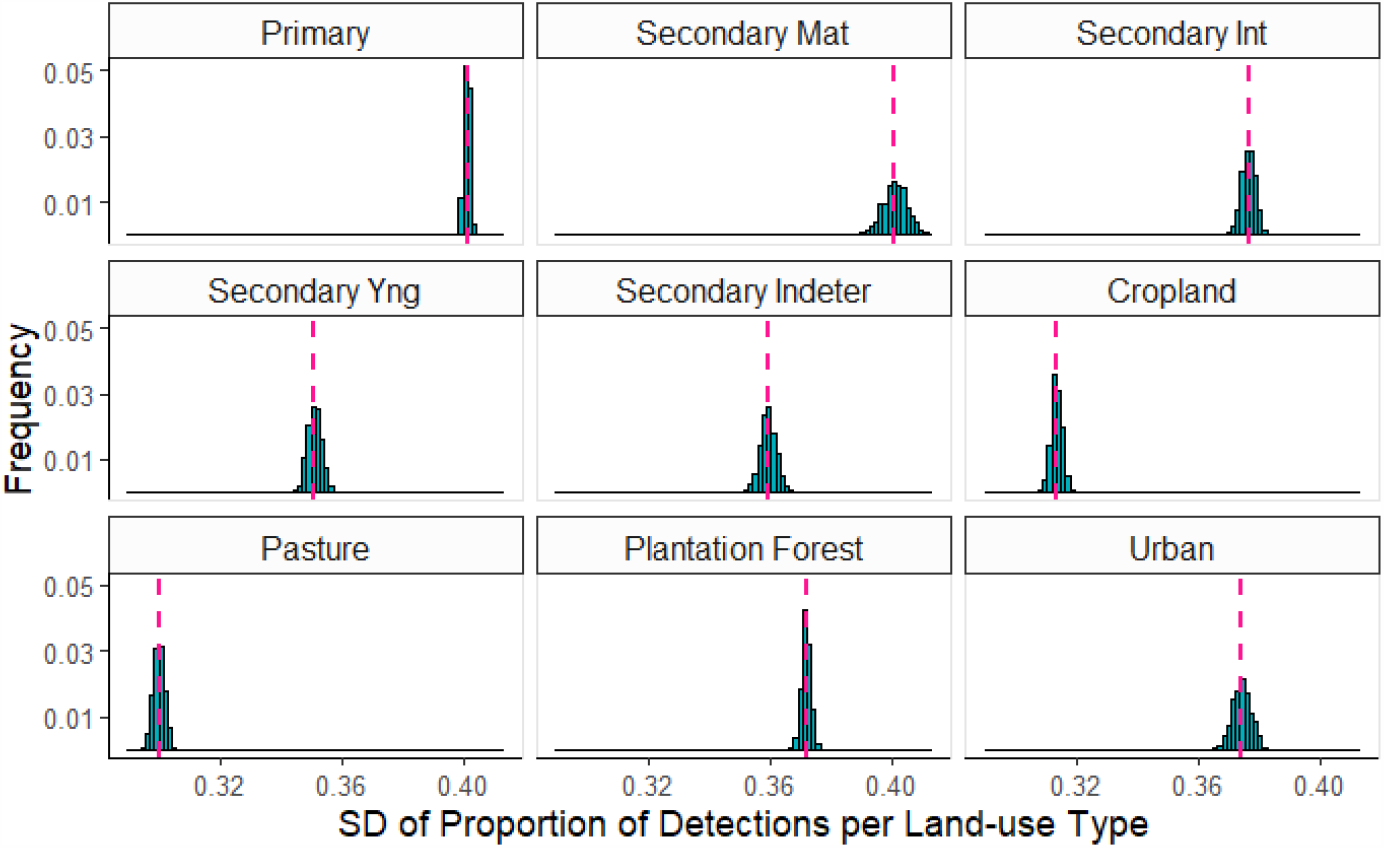
Posterior predictive check for the standard deviation of proportion of species detections observed in each land-cover type. Histograms depict the distribution of test statistics (sd for proportion of detections computed as √ p*1-p) computed for each of 1,000 datasets simulated from the posterior predictive distribution of the Bayesian hierarchical logistic regression model. Vertical dashed lines depict the same summary/test statistic computed from the raw empirical data.

**Fig. S15.**
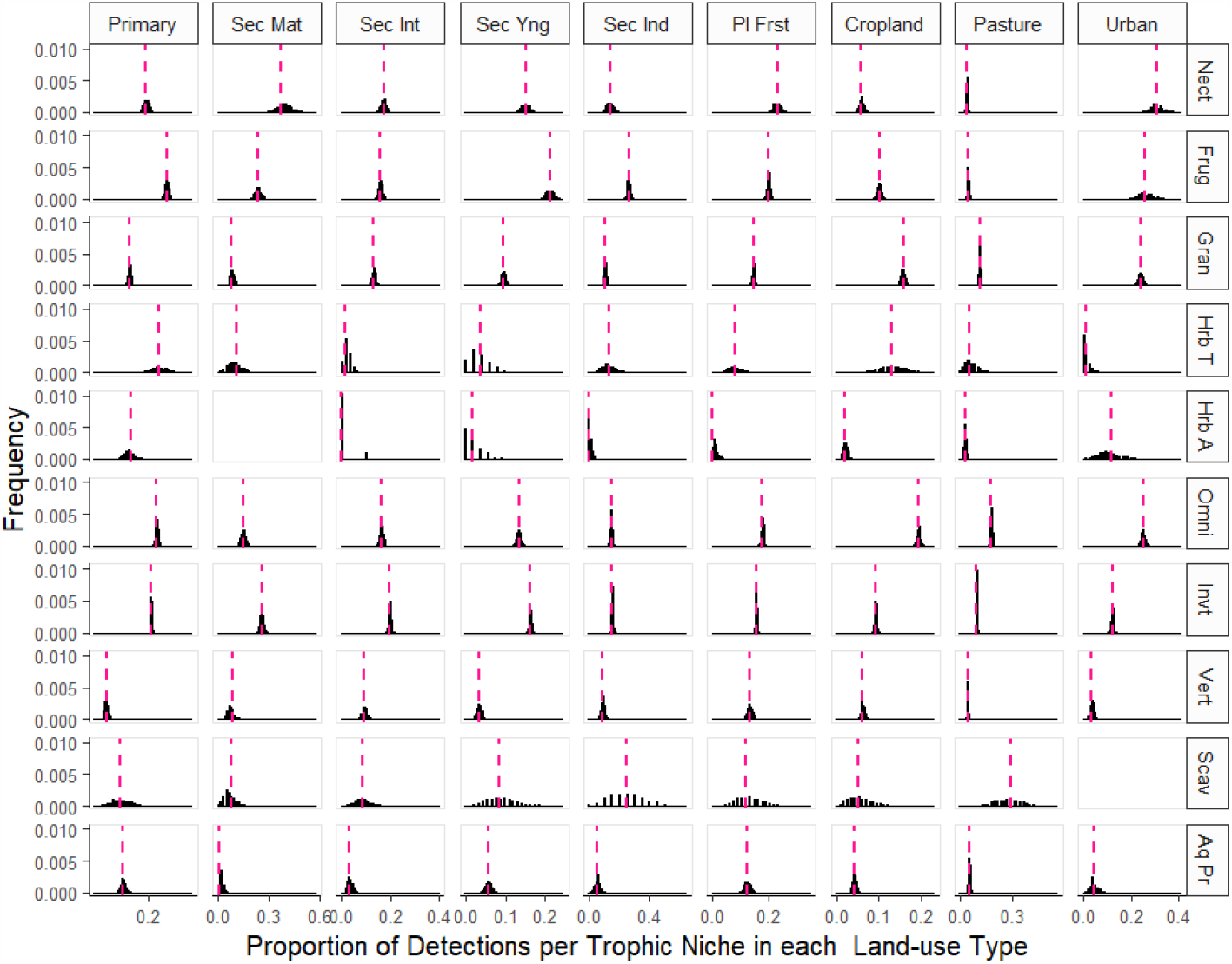
Posterior predictive check for proportion of species detections observed in each land-cover type for each trophic niche. Histograms depict the distribution of test statistics (mean/proportion of detections) computed for each of 1,000 datasets simulated from the posterior predictive distribution of the Bayesian hierarchical logistic regression model. Vertical dashed lines depict the same test statistic computed from the raw empirical data. Abbreviations: Primary (primary vegetation), Sec Mat (seconadry mature vegetation), Sec Int (secondary intermediate vegetation), Sec Ind (secondary vegetation of indeterminate age), Pl First (plantation forest), Nect (nectivore), Frug (frugivore), Gran (granivore), Hrb T (terrestrial herbivore), Hrb A (aquatic herbivore), Omni (omnivore), Invt (invertivore), Vert (vertivore), Scav (scavenger), Aq Pr (aquatic predator).

**Fig. S16.**
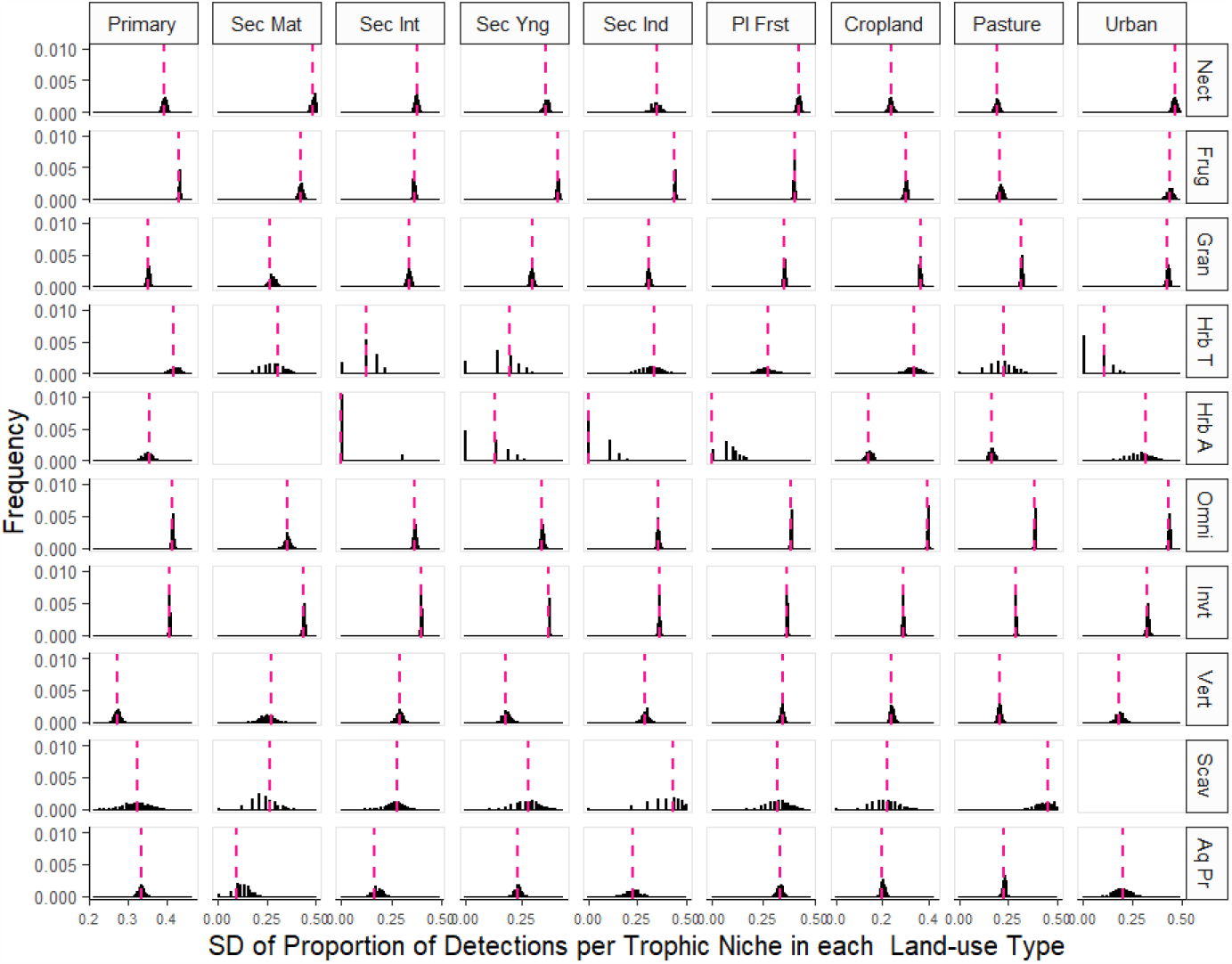
Posterior predictive check for the standard deviation of proportion of species detections observed in each land-cover type for each trophic niche. Histograms depict the distribution of test statistics (sd for proportion of detections computed as √ p*1-p) computed for each of 1,000 datasets simulated from the posterior predictive distribution of the Bayesian hierarchical logistic regression model. Vertical dashed lines depict the same summary/test statistic computed from the raw empirical data. Abbreviations: Primary (primary vegetation), Sec Mat (seconadry mature vegetation), Sec Int (secondary intermediate vegetation), Sec Ind (secondary vegetation of indeterminate age), Pl First (plantation forest), Nect (nectivore), Frug (frugivore), Gran (granivore), Hrb T (terrestrial herbivore), Hrb A (aquatic herbivore), Omni (omnivore), Invt (invertivore), Vert (vertivore), Scav (scavenger), Aq Pr (aquatic predator).

**Fig. S17.**
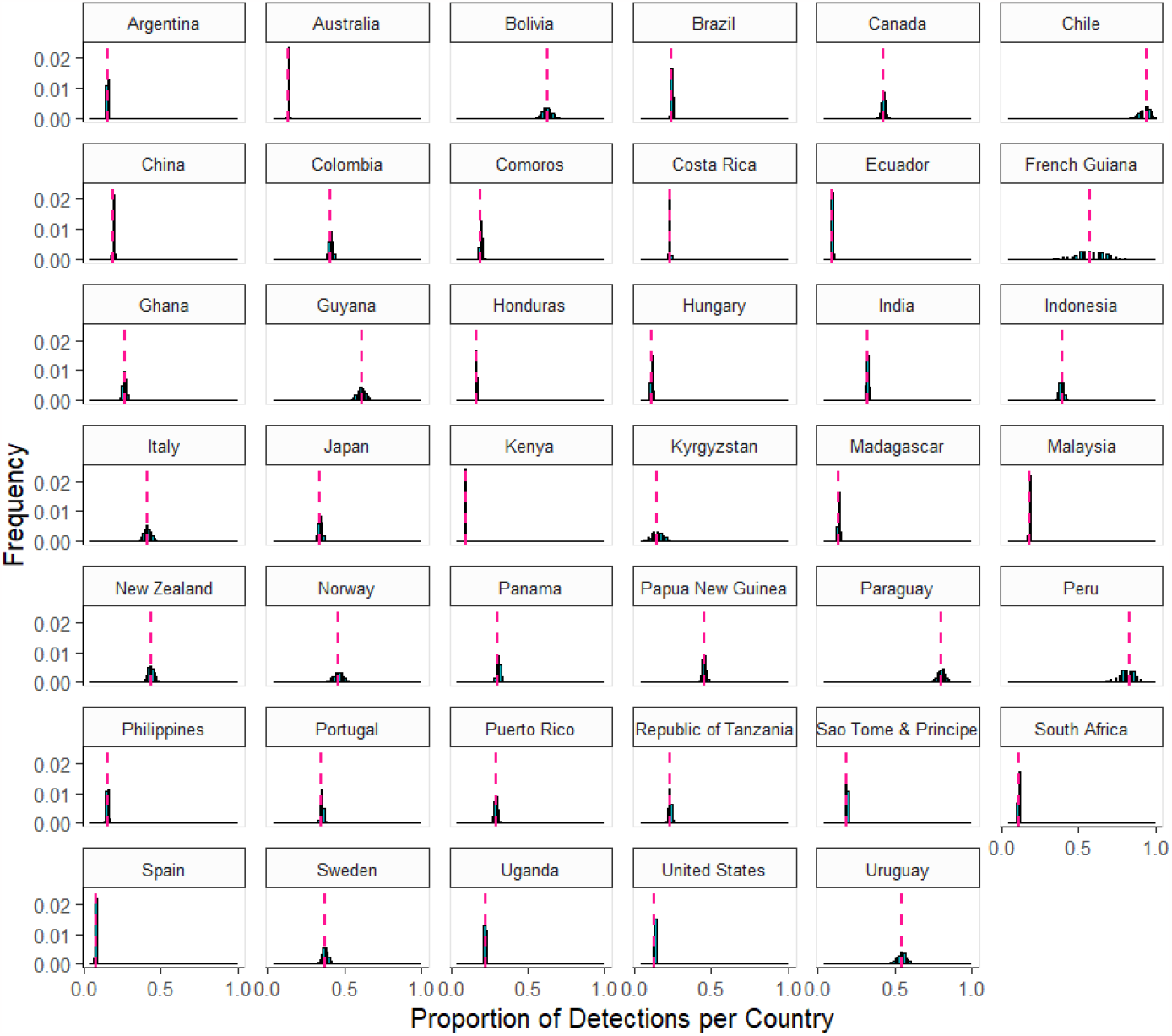
Posterior predictive check for proportion of species detections observed in each country. Histograms depict the distribution of test statistics (mean/proportion of detections) computed for each of 1,000 datasets simulated from the posterior predictive distribution of the Bayesian hierarchical logistic regression model. Vertical dashed lines depict the same test statistic computed from the raw empirical data.

**Fig. S18.**
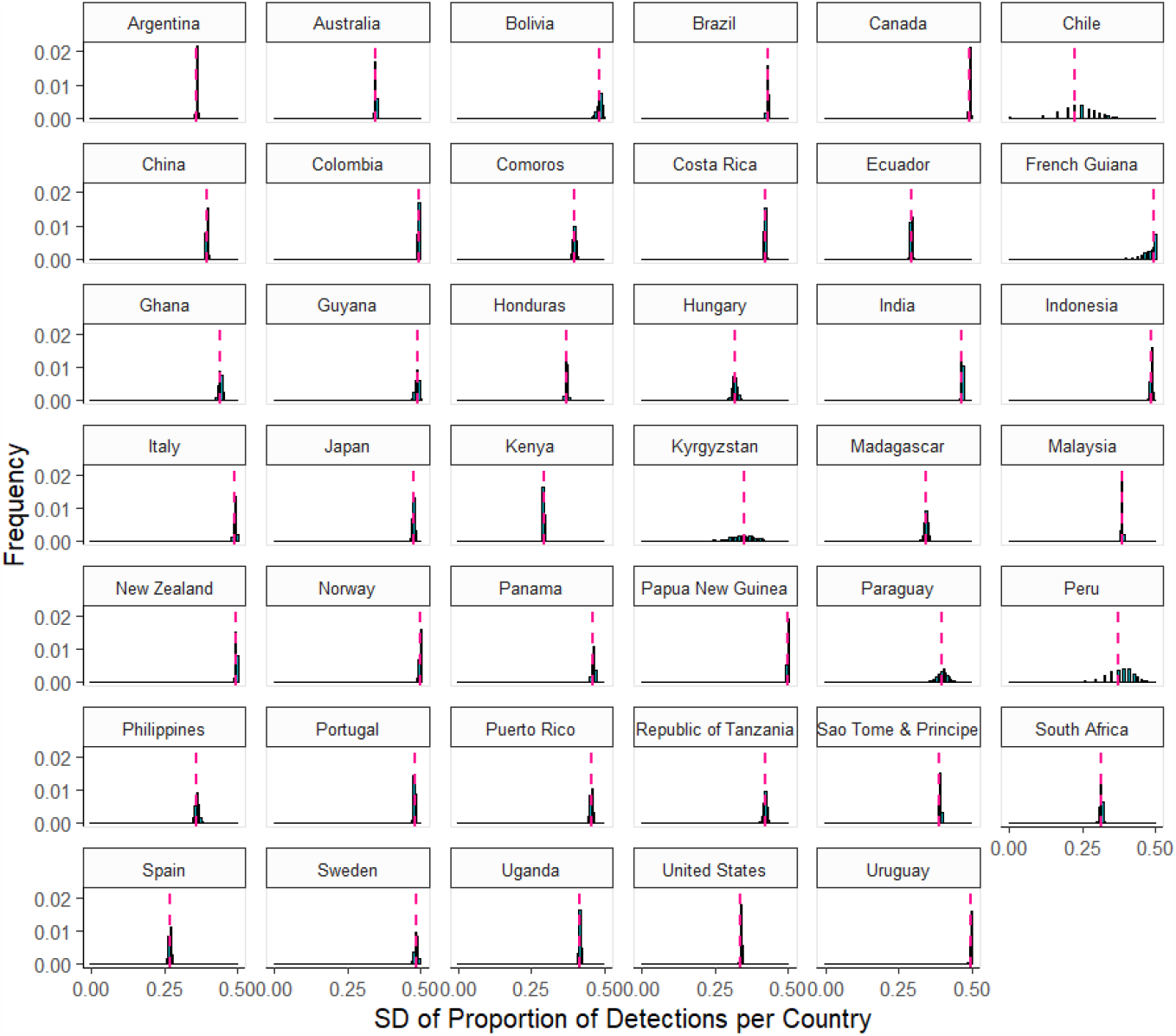
Posterior predictive check for the standard deviation of proportion of species detections observed in each country. Histograms depict the distribution of test statistics (sd for proportion of detections computed as √ p*1-p) computed for each of 1,000 datasets simulated from the posterior predictive distribution of the Bayesian hierarchical logistic regression model. Vertical dashed lines depict the same summary/test statistic computed from the raw empirical data.

**Fig. S19.**
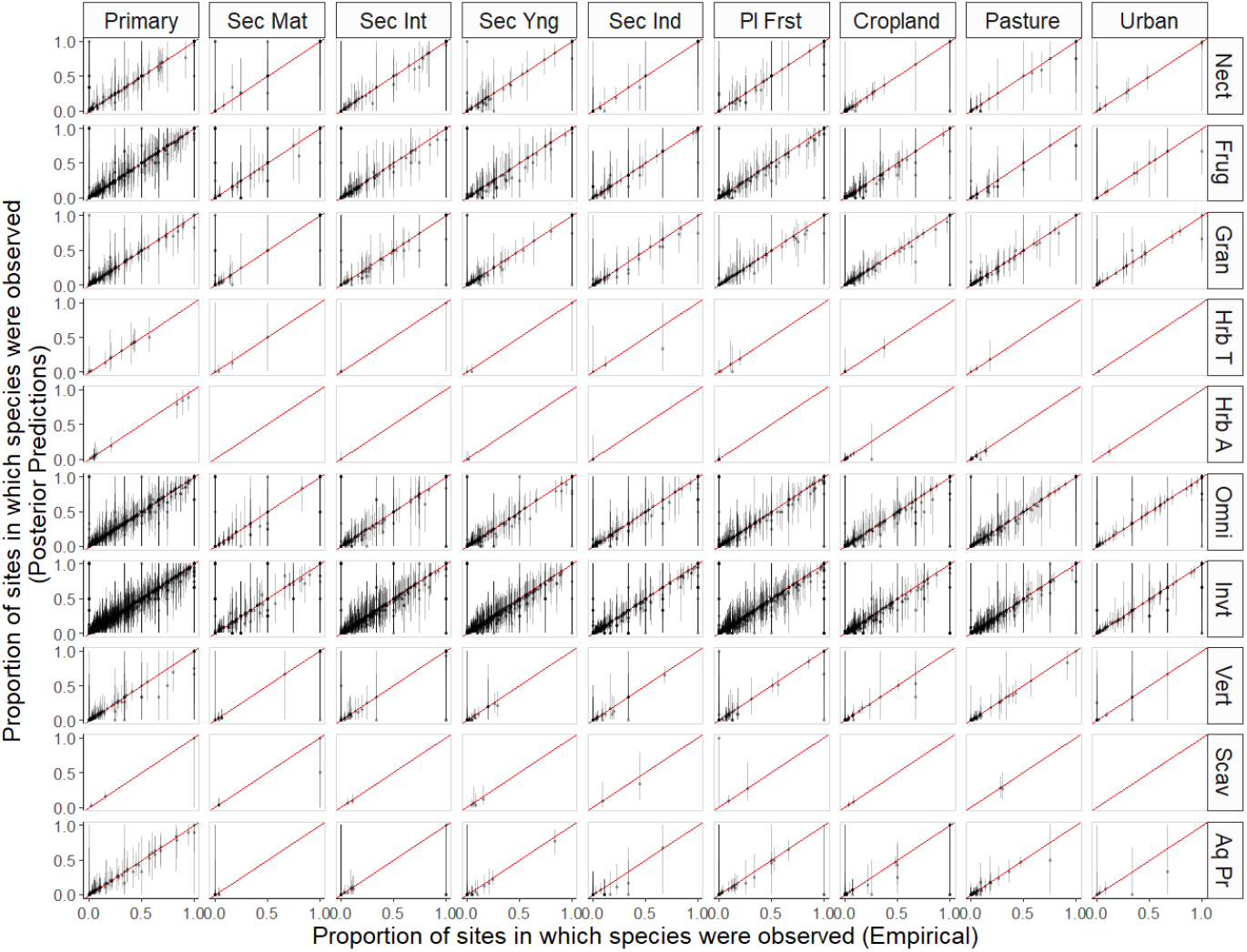
Posterior predictive check for species-level proportions of detections within each land-cover type within which a given species was surveyed for. Y-axes depict distributions of test statistics (mean/proportion of detections per species) computed for each of 1,000 datasets simulated from the posterior predictive distribution of the Bayesian hierarchical logistic regression model. X-axes depict the same test statistic computed for species from the raw empirical data. Diagonal red lines represent the ‘one-to-one’ line along which the distributions of test statistics should fall if the posterior predictive distribution is a good representation of the empirical data. Abbreviations: Primary (primary vegetation), Sec Mat (seconadry mature vegetation), Sec Int (secondary intermediate vegetation), Sec Ind (secondary vegetation of indeterminate age), Pl First (plantation forest), Nect (nectivore), Frug (frugivore), Gran (granivore), Hrb T (terrestrial herbivore), Hrb A (aquatic herbivore), Omni (omnivore), Invt (invertivore), Vert (vertivore), Scav (scavenger), Aq Pr (aquatic predator).

**Fig. S20.**
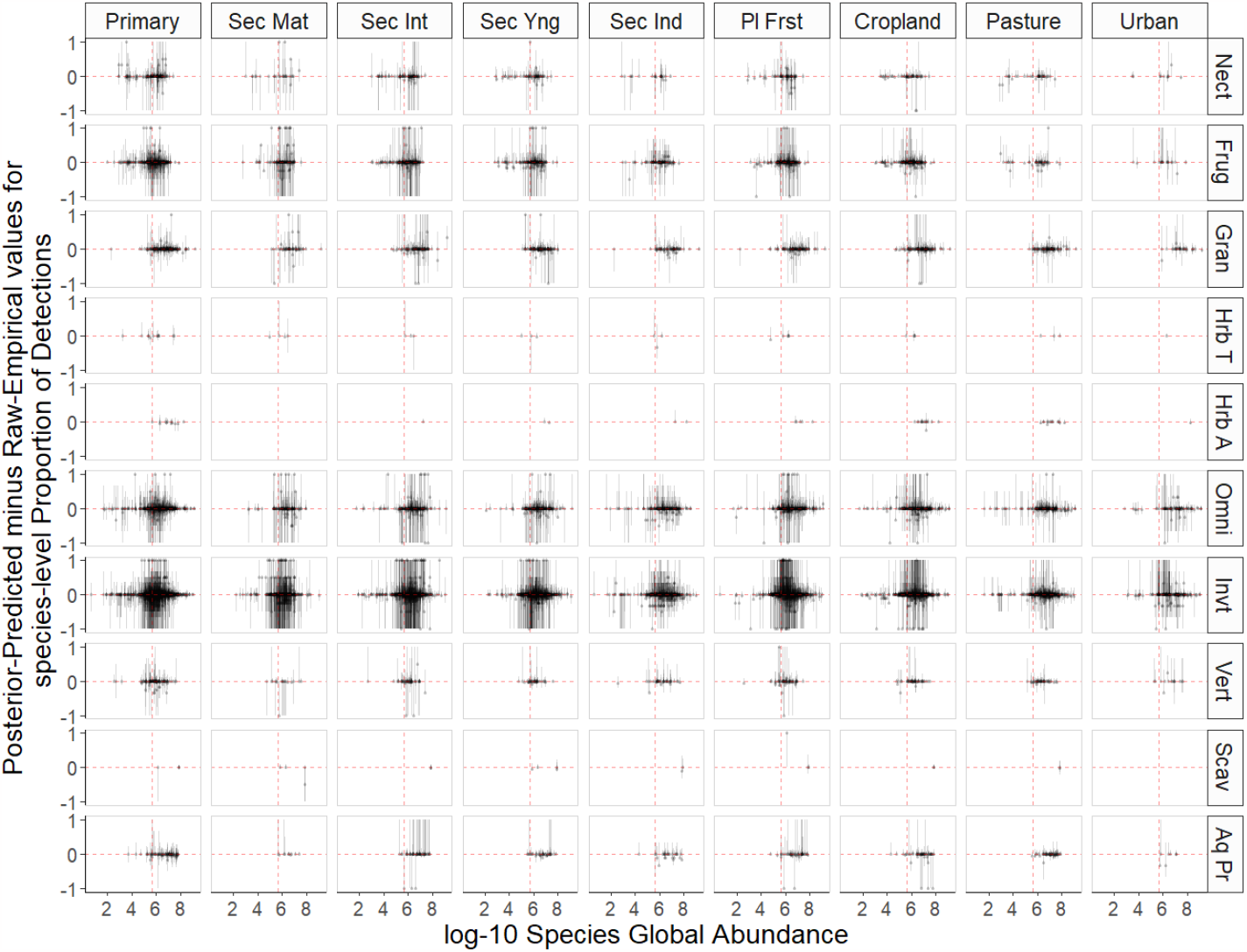
Posterior predictive check for the difference between raw species-level detection proportions and the same quantity computed from posterior predictive simulations (**y-axes**) plotted against species global abundances (**x-axes**). Y-axes depict distributions of test statistics (difference vs empirical data for mean/proportion of detections per species) computed for each of 1,000 datasets simulated from the posterior predictive distribution of the Bayesian hierarchical logistic regression model. X-axes depicts log-10 species global abundnaces. Differences should centre around zero if the posterior predictive distribution is a good representation of the empirical data across the range of species global abundances. Abbreviations: Primary (primary vegetation), Sec Mat (seconadry mature vegetation), Sec Int (secondary intermediate vegetation), Sec Ind (secondary vegetation of indeterminate age), Pl First (plantation forest), Nect (nectivore), Frug (frugivore), Gran (granivore), Hrb T (terrestrial herbivore), Hrb A (aquatic herbivore), Omni (omnivore), Invt (invertivore), Vert (vertivore), Scav (scavenger), Aq Pr (aquatic predator).

**Fig. S21.**
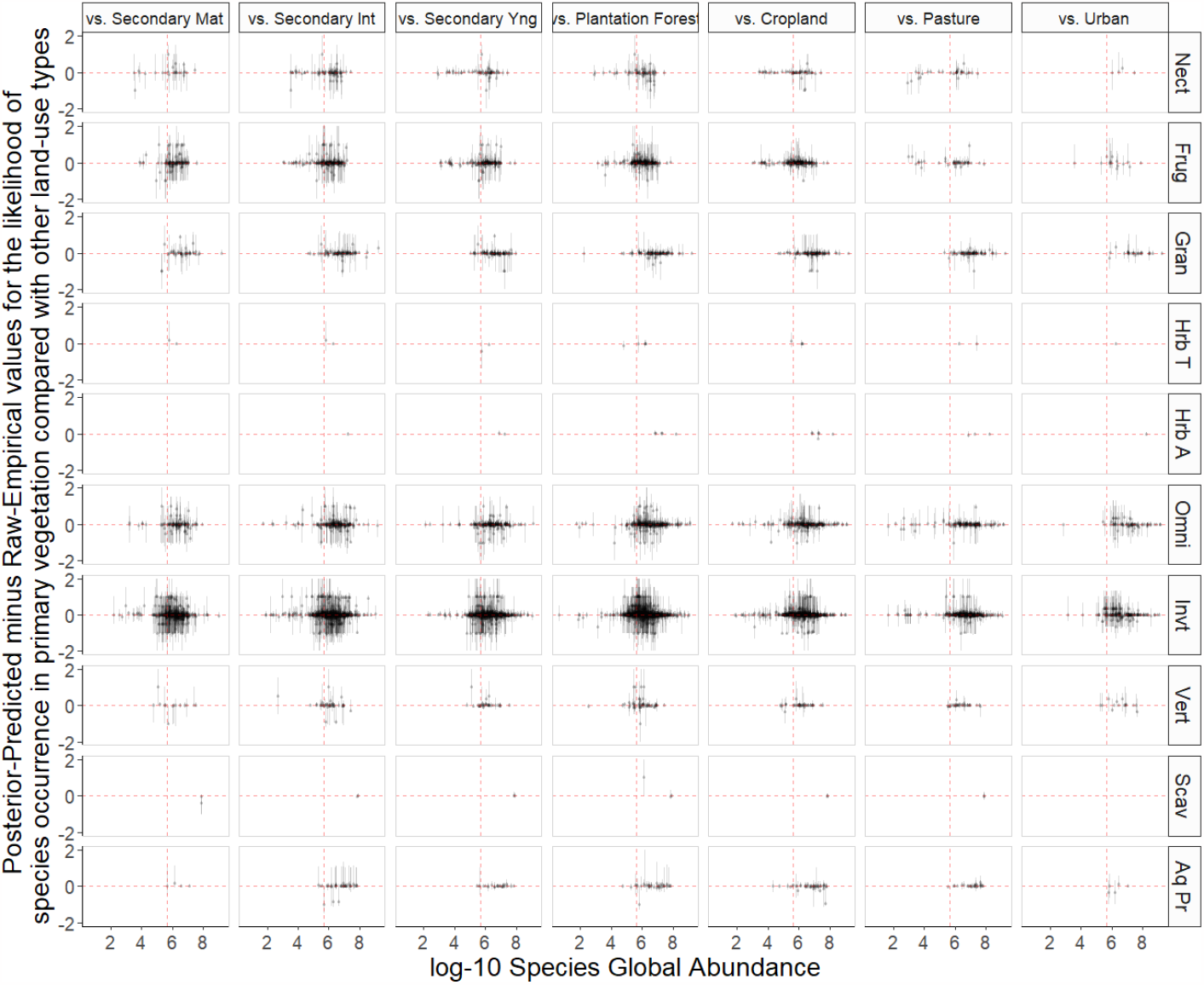
Posterior predictive check representing differences between the empirical likelihood of species occurrence in primary vegetation compared with other land-cover types and the same quantity computed from posterior predictive simulations (**y-axes**) plotted against species global abundances (**x-axes**). Y-axes depict distributions of test statistics (difference vs empirical data for empirical land use contrasts vs primary vegetation) computed for each of 1,000 datasets simulated from the posterior predictive distribution of the Bayesian hierarchical logistic regression model. X-axes depicts log-10 species global abundnaces. Differences should centre around zero if the posterior predictive distribution is a good representation of the empirical data across the range of species global abundances. Abbreviations: Nect (nectivore), Frug (frugivore), Gran (granivore), Hrb T (terrestrial herbivore), Hrb A (aquatic herbivore), Omni (omnivore), Invt (invertivore), Vert (vertivore), Scav (scavenger), Aq Pr (aquatic predator).

**Fig. S22.**
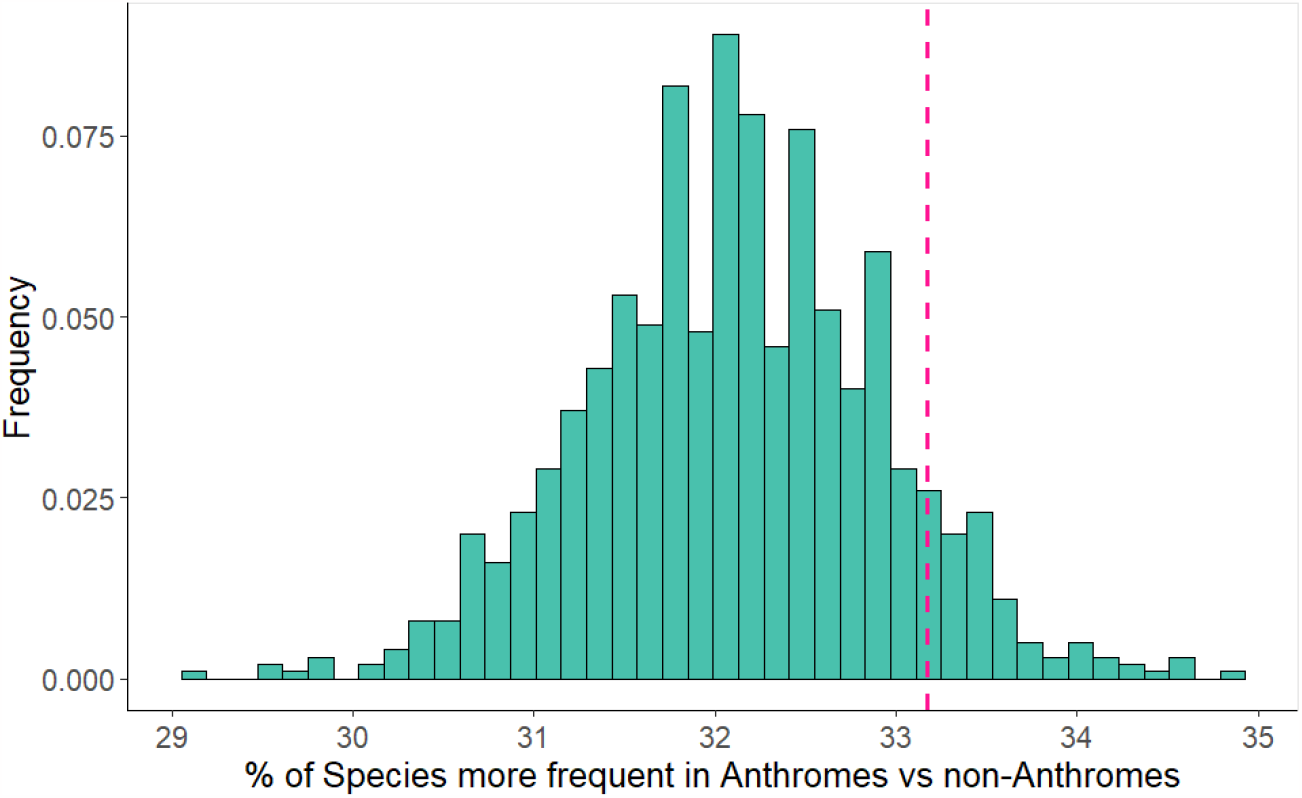
Posterior predictive check for the percentage of species more frequent in anthromes compared with non-anthromes for all bird species recorded in PREDICTS database. The histogram depicts the distribution of test statistics computed for each of 1,000 datasets simulated from the posterior predictive distribution of the Bayesian hierarchical logistic regression model. The vertical dashed line depicts the same test statistic computed from the raw empirical data.

**Fig. S23.**
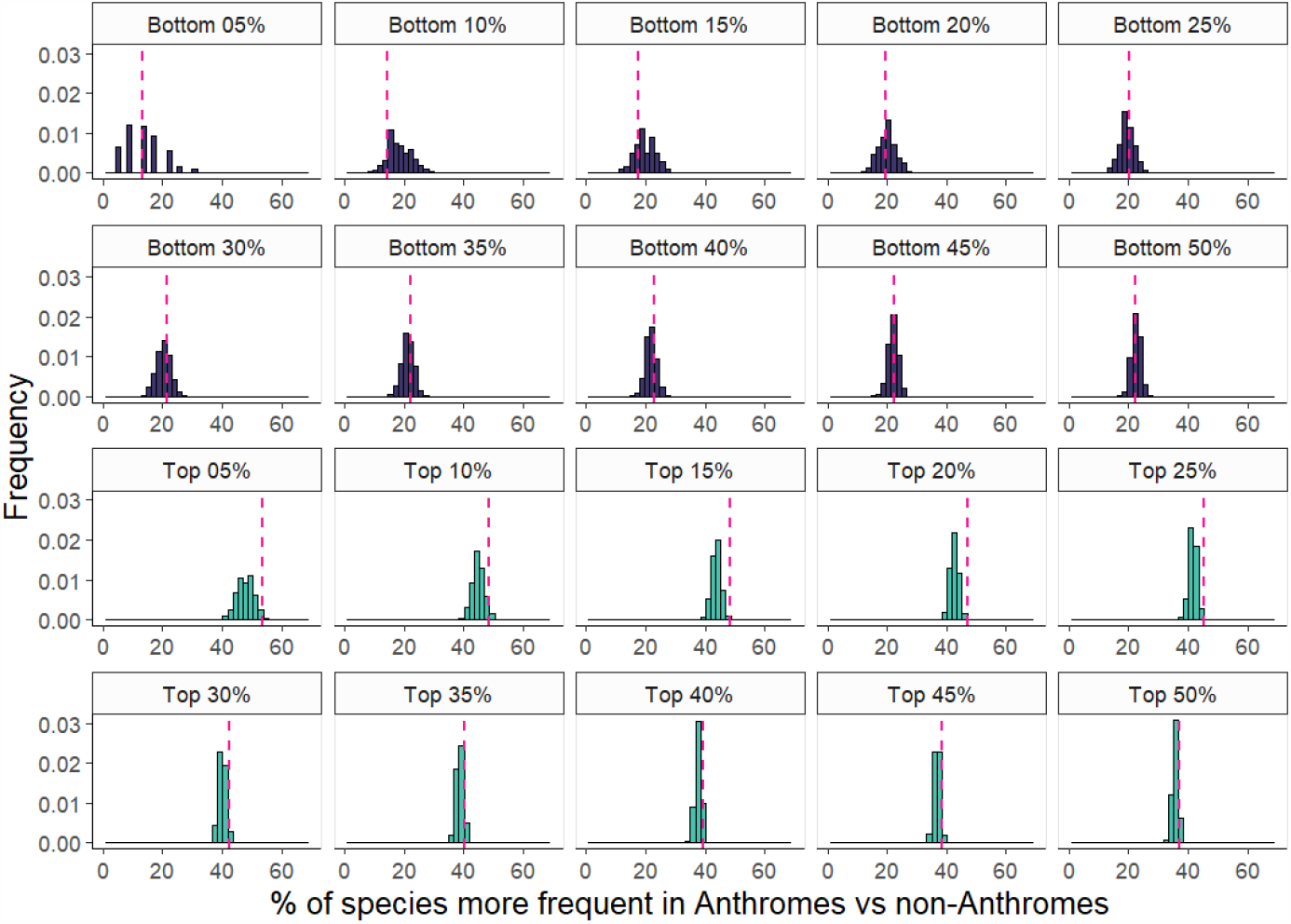
Posterior predictive check for the percentage of species more frequent in anthromes compared with non-anthromes for bird species recorded in PREDICTS database subset by global abundance estimates across *all* bird species. Histograms depict distributions of test statistics computed for each of 1,000 datasets simulated from the posterior predictive distribution of the Bayesian hierarchical logistic regression model. Vertical dashed lines depict the same test statistic computed from the raw empirical data.

**Table S1.**
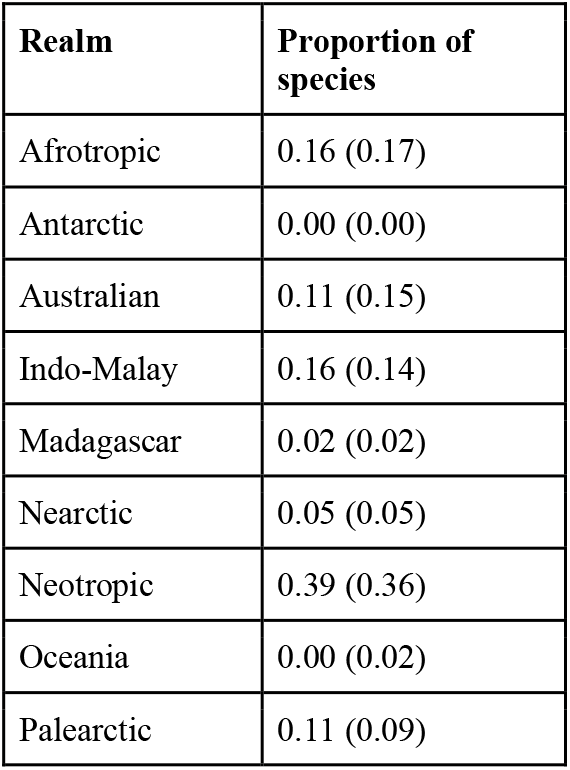
The proportion of species classified as being distributed in each biogeographic realm. Main values correspond to the species included in this study with bracketed values representing the global avifauna. Realm classifications based on (*22*).

| **Realm** | **Proportion of species** |
| --- | --- |
| Afrotropic | 0.16 (0.17) |
| Antarctic | 0.00 (0.00) |
| Australian | 0.11 (0.15) |
| Indo-Malay | 0.16 (0.14) |
| Madagascar | 0.02 (0.02) |
| Nearctic | 0.05 (0.05) |
| Neotropic | 0.39 (0.36) |
| Oceania | 0.00 (0.02) |
| Palearctic | 0.11 (0.09) |

**Table S2.** The proportion of species classified as belonging to each trophic niche. Main values correspond to the species included in this study with bracketed values representing the global avifauna. Trophic niche classifications based on (*22*). Values given to nearest 0.01, with 0.00 indicating less that 0.005 (less than half of one percent of species).

| **Trophic Niche** | **Proportion of species** |
| --- | --- |
| Aquatic predator | 0.04 (0.08) |
| Frugivore | 0.12 (0.10) |
| Granivore | 0.06 (0.07) |
| Herbivore aquatic | 0.01 (0.01) |
| Herbivore terrestrial | 0.00 (0.01) |
| Insectivore | 0.52 (0.48) |
| Nectarivore | 0.04 (0.05) |
| Omnivore | 0.17 (0.17) |
| Scavenger | 0.00 (0.00) |
| Vertivore | 0.04 (0.03) |

**Table S3.**
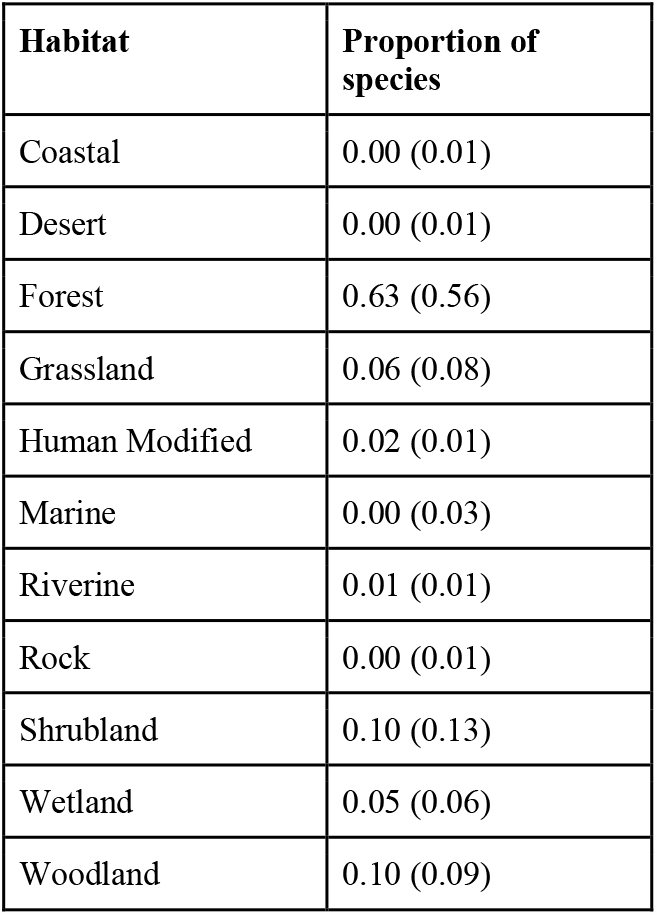
The proportion of species classified as belonging to each habitat type. Main values correspond to the species included in this study with bracketed values representing the global avifauna. Habitat classification based on based on (*23*). Values given to nearest 0.01, with 0.00 indicating less that 0.005 (less than half of one percent of species).

**Table S4.** Summary statistics at the level of land-cover type for models fit to data using global abundance estimates for bird species from Callagahan *et al*. (*19*) and BirdLife International (*24*). Intercepts/Slopes are presented for parameter estimates of scaled-log-global-abundance as an explanatory variable against bird species occurrence. Sample sizes are given for the number of grouping categories at the respective hierarchical level within the model.

| **Land-cover Type** | **Glb Abund Est** | **Intercept Estimate**<br>**( Median (CI) )** | **Slope Estimate**<br>**( Median (CI) )** | **No. Species** | **No. Study** | **No. Site** | **No. Data** |
| --- | --- | --- | --- | --- | --- | --- | --- |
| Primary | Cal | -0.85<br>(-1.49 - -0.12) | -0.11<br>(-0.25 - 0.07) | 2,714 | 74 | 1,912 | 117,774 |
| Secondary Mat | Cal | -1.14<br>(-1.84 - -0.38) | -0.15<br>(-0.4 - 0.07) | 797 | 15 | 86 | 5,720 |
| Secondary Int | Cal | -1.3<br>(-1.92 - -0.53) | -0.16<br>(-0.36 - 0.03) | 1,421 | 26 | 272 | 24,555 |
| Secondary Yng | Cal | -0.97<br>(-1.61 - -0.21) | 0.18<br>(-0.01 - 0.38) | 1,339 | 23 | 374 | 30,966 |
| Secondary Indeter | Cal | -1.21<br>(-1.86 - -0.44) | 0.12<br>(-0.11 - 0.32) | 884 | 16 | 338 | 21,582 |
| Cropland | Cal | -1.16<br>(-1.8 - -0.41) | 0.46<br>(0.28 - 0.64) | 1,374 | 20 | 888 | 65,330. |
| Pasture | Cal | -1.29<br>(-1.95 - -0.53) | 0.22<br>(0.03 - 0.41) | 884 | 18 | 698 | 57,413 |
| Plantation Forest | Cal | -1.08<br>(-1.7 - -0.35) | 0.4<br>(0.22 - 0.58) | 1,713 | 32 | 715 | 59,854 |
| Urban | Cal | -1.85<br>(-2.55 - -1.06) | 0.44<br>(0.2 - 0.71) | 332 | 5 | 171 | 12,111 |
| Primary | BL | -1.11<br>(-1.66 - -0.4) | -0.09<br>(-0.31 - 0.14) | 624 | 68 | 1743 | 33,806 |
| Secondary Mat | BL | -1.34<br>(-2.07 - -0.48) | 0.09<br>(-0.27 - 0.44) | 175 | 13 | 76 | 1,201 |
| Secondary Int | BL | -1.52<br>(-2.1 - -0.76) | -0.12<br>(-0.4 - 0.2) | 378 | 24 | 266 | 8,961 |
| Secondary Yng | BL | -1.18<br>(-1.78 - -0.45) | 0.31<br>(0.03 - 0.61) | 418 | 22 | 372 | 7,746 |
| Secondary Indeter | BL | -1.36<br>(-1.98 - -0.6) | 0.08<br>(-0.24 - 0.38) | 276 | 16 | 338 | 7,776 |
| Cropland | BL | -1.49<br>(-2.07 - -0.73) | 0.6<br>(0.32 - 0.9) | 391 | 20 | 888 | 20,767 |
| Pasture | BL | -1.68<br>(-2.26 - -0.95) | 0.14<br>(-0.14 - 0.43) | 434 | 17 | 697 | 17,171 |
| Plantation Forest | BL | -1.29<br>(-1.83 - -0.54) | 0.31<br>(0.06 - 0.58) | 500 | 31 | 711 | 15,422 |
| Urban | BL | -1.88<br>(-2.59 - -1.03) | 0.36<br>(-0.08 - 0.79) | 100 | 5 | 171 | 6,551 |

**Table S5.** Summary statistics at the level of trophic-niche within land-cover type for models fit to data using bird species global abundance estimates from Callagahan *et al*. (*19*) and BirdLife International (*24*). Intercepts/Slopes are presented for parameter estimates of scaled-log-global-abundance as an explanatory variable against bird species occurrence. Sample sizes are given for the number of grouping categories at the respective hierarchical level within the model.

| Land-cover Type | Trophic-Niche | Glb Abnd Est | Intercept Estimate (Median (CI)) | Slope Estimate (Median (CI)) | No. Species | No. Study | No. Site | No. Data |
| --- | --- | --- | --- | --- | --- | --- | --- | --- |
| Primary | Nectarivore | Cal | -0.83<br>(-1.36 - -0.33) | -0.09<br>(-0.32 - 0.16) | 124 | 43 | 838 | 3,681 |
| Primary | Frugivore | Cal | -0.21<br>(-0.62 - 0.22) | 0<br>(-0.19 - 0.22) | 341 | 64 | 1,525 | 10,574 |
| Primary | Granivore | Cal | -1.31<br>(-1.75 - -0.86) | -0.1<br>(-0.32 - 0.13) | 167 | 64 | 1,665 | 10,682 |
| Primary | Herbivore terrestrial | Cal | -1.6<br>(-2.55 - -0.65) | -0.1<br>(-0.42 - 0.22) | 12 | 14 | 486 | 528 |
| Primary | Herbivore aquatic | Cal | -1.95<br>(-2.86 - -1.03) | -0.07<br>(-0.38 - 0.33) | 12 | 7 | 357 | 492 |
| Primary | Omnivore | Cal | -0.73<br>(-1.1 - -0.35) | -0.2<br>(-0.33 - -0.06) | 449 | 69 | 1741 | 20,640 |
| Primary | Invertivore | Cal | -0.72<br>(-1.07 - -0.38) | -0.24<br>(-0.35 - -0.14) | 1,435 | 66 | 1,684 | 63,347 |
| Primary | Vertivore | Cal | -2.04<br>(-2.57 - -1.51) | -0.1<br>(-0.36 - 0.2) | 96 | 48 | 1,159 | 4,748 |
| Primary | Scavenger | Cal | -1.01<br>(-2.3 - 0.22) | -0.12<br>(-0.42 - 0.22) | 3 | 6 | 116 | 167 |
| Primary | Aquatic predator | Cal | -1.74<br>(-2.27 - -1.21) | -0.09<br>(-0.37 - 0.23) | 75 | 29 | 760 | 2,915 |
| Secondary Mat | Nectarivore | Cal | -0.94<br>(-1.69 - -0.15) | -0.15<br>(-0.48 - 0.16) | 28 | 10 | 38 | 83 |
| Secondary Mat | Frugivore | Cal | -0.82<br>(-1.37 - -0.27) | -0.07<br>(-0.4 - 0.23) | 119 | 14 | 84 | 684 |
| Secondary Mat | Granivore | Cal | -1.58<br>(-2.24 - -0.97) | -0.21<br>(-0.59 - 0.09) | 52 | 12 | 75 | 590 |
| Secondary Mat | Herbivore terrestrial | Cal | -1.65<br>(-2.74 - -0.57) | -0.16<br>(-0.58 - 0.2) | 4 | 3 | 34 | 65 |
| Secondary Mat | Omnivore | Cal | -1.13<br>(-1.67 - -0.6) | -0.19<br>(-0.48 - 0.08) | 110 | 14 | 78 | 887 |
| Secondary Mat | Invertivore | Cal | -0.76<br>(-1.19 - -0.33) | -0.13<br>(-0.33 - 0.07) | 452 | 14 | 78 | 2,895 |
| Secondary Mat | Vertivore | Cal | -1.95<br>(-2.72 - -1.19) | -0.15<br>(-0.53 - 0.19) | 18 | 8 | 66 | 230 |
| Secondary Mat | Scavenger | Cal | -1.21<br>(-2.5 - 0.04) | -0.12<br>(-0.51 - 0.26) | 4 | 3 | 33 | 65 |
| Secondary Mat | Aquatic predator | Cal | -2.64<br>(-3.51 - -1.82) | -0.18<br>(-0.59 - 0.17) | 10 | 4 | 34 | 221 |
| Secondary Int | Nectarivore | Cal | -0.74<br>(-1.31 - -0.2) | -0.1<br>(-0.36 - 0.18) | 75 | 16 | 162 | 1,463 |
| Secondary Int | Frugivore | Cal | -0.86<br>(-1.33 - -0.38) | -0.1<br>(-0.34 - 0.15) | 185 | 22 | 232 | 3,388 |
| Secondary Int | Granivore | Cal | -1.61<br>(-2.12 - -1.06) | -0.26<br>(-0.59 - -0.01) | 95 | 23 | 245 | 1,993 |
| Secondary Int | Herbivore terrestrial | Cal | -1.91<br>(-3.01 - -0.86) | -0.15<br>(-0.52 - 0.2) | 3 | 3 | 62 | 62 |
| Secondary Int | Herbivore aquatic | Cal | -2.63<br>(-3.76 - -1.48) | -0.15<br>(-0.53 - 0.24) | 1 | 1 | 10 | 10 |
| Secondary Int | Omnivore | Cal | -1.1<br>(-1.52 - -0.62) | -0.18<br>(-0.38 - 0.01) | 211 | 24 | 266 | 3,586 |
| Secondary Int | Invertivore | Cal | -0.94<br>(-1.3 - -0.55) | -0.12<br>(-0.26 - 0.02) | 752 | 24 | 266 | 12,538 |
| Secondary Int | Vertivore | Cal | -2.47<br>(-3.11 - -1.85) | -0.13<br>(-0.43 - 0.2) | 48 | 17 | 189 | 755 |
| Secondary Int | Scavenger | Cal | -1.39<br>(-2.74 - -0.14) | -0.15<br>(-0.48 - 0.22) | 2 | 4 | 77 | 151 |
| Secondary Int | Aquatic predator | Cal | -2.93<br>(-3.64 - -2.23) | -0.2<br>(-0.6 - 0.14) | 49 | 14 | 186 | 609 |
| Secondary Yng | Nectarivore | Cal | -0.54<br>(-1.12 - 0.06) | 0.2<br>(-0.06 - 0.49) | 63 | 16 | 280 | 1,249 |
| Secondary Yng | Frugivore | Cal | -0.53<br>(-1.01 - -0.05) | 0.28<br>(0.06 - 0.56) | 154 | 19 | 312 | 2,291 |
| Secondary Yng | Granivore | Cal | -1.09<br>(-1.58 - -0.58) | 0.22<br>(-0.06 - 0.53) | 103 | 21 | 368 | 3,552 |
| Secondary Yng | Herbivore terrestrial | Cal | -1.54<br>(-2.68 - -0.42) | 0.18<br>(-0.16 - 0.55) | 3 | 2 | 26 | 51 |
| Secondary Yng | Herbivore aquatic | Cal | -2.34<br>(-3.38 - -1.25) | 0.18<br>(-0.17 - 0.56) | 2 | 2 | 55 | 55 |
| Secondary Yng | Omnivore | Cal | -0.85<br>(-1.26 - -0.41) | 0.11<br>(-0.11 - 0.32) | 202 | 23 | 374 | 5,005 |
| Secondary Yng | Invertivore | Cal | -0.88<br>(-1.23 - -0.51) | 0.08<br>(-0.09 - 0.24) | 734 | 23 | 374 | 16,467 |
| Secondary Yng | Vertivore | Cal | -2.23<br>(-2.9 - -1.55) | 0.17<br>(-0.17 - 0.51) | 37 | 14 | 273 | 1,252 |
| Secondary Yng | Scavenger | Cal | -1.14<br>(-2.41 - 0.05) | 0.16<br>(-0.17 - 0.51) | 4 | 5 | 51 | 93 |
| Secondary Yng | Aquatic predator | Cal | -2.31<br>(-2.98 - -1.63) | 0.16<br>(-0.19 - 0.5) | 37 | 13 | 222 | 951 |
| Secondary Indeter | Nectarivore | Cal | -0.65<br>(-1.34 - 0.11) | 0.13<br>(-0.19 - 0.43) | 23 | 9 | 101 | 257 |
| Secondary Indeter | Frugivore | Cal | -0.76<br>(-1.26 - -0.23) | 0.15<br>(-0.18 - 0.44) | 89 | 13 | 198 | 1,741 |
| Secondary Indeter | Granivore | Cal | -1.27<br>(-1.81 - -0.73) | 0.08<br>(-0.23 - 0.33) | 64 | 14 | 275 | 2,263 |
| Secondary Indeter | Herbivore terrestrial | Cal | -1.76<br>(-2.87 - -0.64) | 0.13<br>(-0.27 - 0.46) | 3 | 3 | 83 | 83 |
| Secondary Indeter | Herbivore aquatic | Cal | -2.62<br>(-3.75 - -1.52) | 0.11<br>(-0.29 - 0.5) | 2 | 2 | 80 | 80 |
| Secondary Indeter | Omnivore | Cal | -1.21<br>(-1.65 - -0.75) | 0.13<br>(-0.07 - 0.32) | 178 | 16 | 338 | 4,526 |
| Secondary Indeter | Invertivore | Cal | -1.33<br>(-1.71 - -0.93) | 0.16<br>(0.01 - 0.33) | 456 | 16 | 338 | 11,429 |
| Secondary Indeter | Vertivore | Cal | -2.47<br>(-3.16 - -1.8) | 0.07<br>(-0.29 - 0.38) | 40 | 14 | 298 | 870 |
| Secondary Indeter | Scavenger | Cal | -1.33<br>(-2.68 - -0.04) | 0.13<br>(-0.28 - 0.47) | 2 | 2 | 11 | 20 |
| Secondary Indeter | Aquatic predator | Cal | -2.47<br>(-3.16 - -1.79) | 0.11<br>(-0.26 - 0.44) | 27 | 8 | 107 | 313 |
| Cropland | Nectarivore | Cal | -0.79<br>(-1.4 - -0.19) | 0.48<br>(0.19 - 0.76) | 53 | 12 | 525 | 2,955 |
| Cropland | Frugivore | Cal | -0.98<br>(-1.44 - -0.52) | 0.51<br>(0.27 - 0.76) | 169 | 16 | 628 | 5,243 |
| Cropland | Granivore | Cal | -0.52<br>(-0.98 - -0.04) | 0.46<br>(0.19 - 0.72) | 112 | 18 | 683 | 8,319 |
| Cropland | Herbivore terrestrial | Cal | -1.75<br>(-2.78 - -0.66) | 0.46<br>(0.11 - 0.82) | 4 | 4 | 153 | 153 |
| Cropland | Herbivore aquatic | Cal | -2.69<br>(-3.6 - -1.74) | 0.45<br>(0.13 - 0.83) | 15 | 7 | 298 | 1,764 |
| Cropland | Omnivore | Cal | -0.74<br>(-1.15 - -0.3) | 0.48<br>(0.3 - 0.66) | 237 | 20 | 888 | 10,794 |
| Cropland | Invertivore | Cal | -1.57<br>(-1.93 - -1.19) | 0.4<br>(0.26 - 0.55) | 683 | 20 | 888 | 29,388 |
| Cropland | Vertivore | Cal | -2.3<br>(-2.92 - -1.67) | 0.44<br>(0.13 - 0.76) | 47 | 17 | 796 | 3,660 |
| Cropland | Scavenger | Cal | -1.35<br>(-2.76 - -0.12) | 0.45<br>(0.11 - 0.8) | 2 | 2 | 64 | 117 |
| Cropland | Aquatic predator | Cal | -2.58<br>(-3.22 - -1.93) | 0.44<br>(0.11 - 0.76) | 52 | 14 | 612 | 2,937 |
| Pasture | Nectarivore | Cal | -1.19<br>(-1.85 - -0.53) | 0.17<br>(-0.13 - 0.46) | 38 | 8 | 568 | 2,247 |
| Pasture | Frugivore | Cal | -1.29<br>(-1.86 - -0.7) | 0.33<br>(0.08 - 0.63) | 48 | 9 | 587 | 2,110 |
| Pasture | Granivore | Cal | -1.16<br>(-1.62 - -0.68) | 0.19<br>(-0.1 - 0.42) | 89 | 16 | 665 | 8,429 |
| Pasture | Herbivore terrestrial | Cal | -1.97<br>(-3 - -0.88) | 0.2<br>(-0.14 - 0.56) | 3 | 3 | 72 | 72 |
| Pasture | Herbivore aquatic | Cal | -2.67<br>(-3.6 - -1.75) | 0.22<br>(-0.15 - 0.61) | 14 | 5 | 271 | 1,795 |
| Pasture | Omnivore | Cal | -1.1<br>(-1.52 - -0.68) | 0.21<br>(0.01 - 0.41) | 178 | 18 | 698 | 10,126 |
| Pasture | Invertivore | Cal | -1.49<br>(-1.87 - -1.09) | 0.2<br>(0.04 - 0.36) | 422 | 18 | 698 | 25,198 |
| Pasture | Vertivore | Cal | -2.21<br>(-2.82 - -1.6) | 0.2<br>(-0.13 - 0.53) | 41 | 15 | 693 | 3,663 |
| Pasture | Scavenger | Cal | -1.33<br>(-2.62 - -0.05) | 0.24<br>(-0.11 - 0.58) | 2 | 3 | 36 | 56 |
| Pasture | Aquatic predator | Cal | -2.27<br>(-2.86 - -1.64) | 0.22<br>(-0.11 - 0.56) | 49 | 13 | 657 | 3,717 |
| Plantation Forest | Nectarivore | Cal | -0.22<br>(-0.78 - 0.41) | 0.45<br>(0.2 - 0.76) | 63 | 17 | 325 | 930 |
| Plantation Forest | Frugivore | Cal | -1.04<br>(-1.49 - -0.57) | 0.54<br>(0.31 - 0.81) | 210 | 25 | 569 | 7,667 |
| Plantation Forest | Granivore | Cal | -0.97<br>(-1.44 - -0.47) | 0.31<br>(0.03 - 0.54) | 97 | 30 | 697 | 5,556 |
| Plantation Forest | Herbivore terrestrial | Cal | -1.61<br>(-2.69 - -0.56) | 0.39<br>(0.06 - 0.74) | 5 | 5 | 238 | 238 |
| Plantation Forest | Herbivore aquatic | Cal | -2.69<br>(-3.64 - -1.62) | 0.37<br>(-0.02 - 0.73) | 5 | 4 | 134 | 216 |
| Plantation Forest | Omnivore | Cal | -1.14<br>(-1.53 - -0.74) | 0.34<br>(0.16 - 0.52) | 283 | 32 | 715 | 10,339 |
| Plantation Forest | Invertivore | Cal | -1.38<br>(-1.73 - -1) | 0.39<br>(0.26 - 0.53) | 946 | 32 | 715 | 32,074 |
| Plantation Forest | Vertivore | Cal | -1.94<br>(-2.54 - -1.32) | 0.4<br>(0.09 - 0.72) | 56 | 22 | 453 | 1,651 |
| Plantation Forest | Scavenger | Cal | -1.31<br>(-2.72 - -0.02) | 0.39<br>(0.05 - 0.74) | 3 | 4 | 56 | 68 |
| Plantation Forest | Aquatic predator | Cal | -2.23<br>(-2.85 - -1.57) | 0.38<br>(0.06 - 0.7) | 45 | 17 | 291 | 1,115 |
| Urban | Nectarivore | Cal | -1.41<br>(-2.25 - -0.59) | 0.49<br>(0.18 - 0.88) | 10 | 4 | 84 | 285 |
| Urban | Frugivore | Cal | -1.58<br>(-2.32 - -0.83) | 0.51<br>(0.19 - 0.87) | 19 | 5 | 171 | 278 |
| Urban | Granivore | Cal | -2.01<br>(-2.73 - -1.33) | 0.49<br>(0.16 - 0.91) | 30 | 5 | 171 | 1,573 |
| Urban | Herbivore terrestrial | Cal | -2.44<br>(-3.57 - -1.25) | 0.44<br>(0.09 - 0.84) | 1 | 1 | 87 | 87 |
| Urban | Herbivore aquatic | Cal | -3.09<br>(-4.26 - -1.93) | 0.47<br>(0.1 - 0.92) | 1 | 1 | 87 | 87 |
| Urban | Omnivore | Cal | -1.38<br>(-1.94 - -0.78) | 0.41<br>(0.17 - 0.68) | 81 | 5 | 171 | 3,031 |
| Urban | Invertivore | Cal | -2.13<br>(-2.66 - -1.63) | 0.28<br>(0.04 - 0.51) | 168 | 5 | 171 | 5,788 |
| Urban | Vertivore | Cal | -3.1<br>(-3.93 - -2.3) | 0.43<br>(0.07 - 0.83) | 15 | 4 | 131 | 765 |
| Urban | Aquatic predator | Cal | -3.14<br>(-4.02 - -2.25) | 0.42<br>(0.05 - 0.8) | 7 | 4 | 131 | 217 |
| Primary | Nectarivore | BL | -1.28<br>(-1.99 - -0.56) | -0.08<br>(-0.46 - 0.35) | 26 | 11 | 166 | 875 |
| Primary | Frugivore | BL | -0.71<br>(-1.31 - -0.12) | -0.08<br>(-0.33 - 0.25) | 60 | 32 | 811 | 2,096 |
| Primary | Granivore | BL | -1.36<br>(-1.98 - -0.74) | -0.07<br>(-0.39 - 0.33) | 37 | 31 | 869 | 2,764 |
| Primary | Herbivore terrestrial | BL | -1.42<br>(-2.54 - -0.38) | -0.11<br>(-0.56 - 0.34) | 6 | 5 | 273 | 315 |
| Primary | Herbivore aquatic | BL | -1.8<br>(-2.72 - -0.88) | -0.02<br>(-0.38 - 0.62) | 10 | 6 | 333 | 430 |
| Primary | Omnivore | BL | -1.06<br>(-1.51 - -0.59) | -0.2<br>(-0.47 - 0.04) | 119 | 51 | 1,342 | 6,865 |
| Primary | Invertivore | BL | -1.09<br>(-1.49 - -0.67) | -0.16<br>(-0.36 - 0.03) | 283 | 55 | 1,591 | 17,750 |
| Primary | Vertivore | BL | -1.97<br>(-2.65 - -1.23) | 0.04<br>(-0.25 - 0.48) | 41 | 35 | 813 | 1,286 |
| Primary | Scavenger | BL | -1.27<br>(-2.65 - 0.13) | -0.09<br>(-0.5 - 0.39) | 1 | 1 | 1 | 1 |
| Primary | Aquatic predator | BL | -1.91<br>(-2.54 - -1.21) | -0.18<br>(-0.65 - 0.14) | 41 | 18 | 474 | 1,424 |
| Secondary Mat | Nectarivore | BL | -1.35<br>(-2.3 - -0.44) | 0.07<br>(-0.45 - 0.54) | 8 | 4 | 7 | 20 |
| Secondary Mat | Frugivore | BL | -1.09<br>(-1.91 - -0.26) | 0.11<br>(-0.32 - 0.54) | 26 | 6 | 33 | 124 |
| Secondary Mat | Granivore | BL | -1.63<br>(-2.57 - -0.79) | 0.08<br>(-0.42 - 0.57) | 11 | 6 | 42 | 81 |
| Secondary Mat | Omnivore | BL | -1.04<br>(-1.78 - -0.29) | 0.02<br>(-0.41 - 0.42) | 26 | 11 | 69 | 127 |
| Secondary Mat | Invertivore | BL | -1.33<br>(-1.93 - -0.71) | 0.08<br>(-0.29 - 0.43) | 86 | 10 | 73 | 552 |
| Secondary Mat | Vertivore | BL | -2.08<br>(-3.05 - -1.2) | 0.15<br>(-0.31 - 0.67) | 11 | 5 | 40 | 109 |
| Secondary Mat | Scavenger | BL | -1.55<br>(-2.98 - -0.08) | 0.12<br>(-0.37 - 0.65) | 2 | 1 | 31 | 62 |
| Secondary Mat | Aquatic predator | BL | -2.45<br>(-3.38 - -1.56) | 0.07<br>(-0.44 - 0.6) | 5 | 3 | 33 | 126 |
| Secondary Int | Nectarivore | BL | -1.45<br>(-2.24 - -0.69) | -0.13<br>(-0.54 - 0.3) | 17 | 9 | 102 | 460 |
| Secondary Int | Frugivore | BL | -1.17<br>(-1.76 - -0.53) | -0.08<br>(-0.4 - 0.32) | 38 | 13 | 193 | 1,054 |
| Secondary Int | Granivore | BL | -1.66<br>(-2.4 - -0.97) | -0.15<br>(-0.59 - 0.27) | 23 | 13 | 163 | 475 |
| Secondary Int | Herbivore aquatic | BL | -2.21<br>(-3.3 - -1.22) | -0.07<br>(-0.51 - 0.64) | 1 | 1 | 10 | 10 |
| Secondary Int | Omnivore | BL | -1.41<br>(-1.93 - -0.89) | -0.15<br>(-0.47 - 0.17) | 63 | 22 | 263 | 1,466 |
| Secondary Int | Invertivore | BL | -1.25<br>(-1.69 - -0.78) | -0.19<br>(-0.45 - 0.07) | 179 | 22 | 250 | 4,696 |
| Secondary Int | Vertivore | BL | -2.52<br>(-3.27 - -1.78) | -0.03<br>(-0.42 - 0.46) | 26 | 14 | 159 | 425 |
| Secondary Int | Aquatic predator | BL | -2.77<br>(-3.63 - -1.99) | -0.11<br>(-0.57 - 0.42) | 31 | 11 | 169 | 375 |
| Secondary Yng | Nectarivore | BL | -1.02<br>(-1.75 - -0.26) | 0.28<br>(-0.16 - 0.68) | 23 | 7 | 42 | 339 |
| Secondary Yng | Frugivore | BL | -0.83<br>(-1.46 - -0.2) | 0.35<br>(0.01 - 0.74) | 46 | 11 | 105 | 635 |
| Secondary Yng | Granivore | BL | -1.28<br>(-1.99 - -0.57) | 0.37<br>(-0.01 - 0.86) | 25 | 12 | 244 | 489 |
| Secondary Yng | Herbivore aquatic | BL | -1.86<br>(-2.95 - -0.8) | 0.36<br>(-0.1 - 1.09) | 1 | 1 | 47 | 47 |
| Secondary Yng | Omnivore | BL | -1.03<br>(-1.54 - -0.5) | 0.22<br>(-0.15 - 0.54) | 72 | 21 | 352 | 1,340 |
| Secondary Yng | Invertivore | BL | -1.21<br>(-1.65 - -0.77) | 0.29<br>(0.02 - 0.54) | 208 | 21 | 371 | 3,831 |
| Secondary Yng | Vertivore | BL | -2.21<br>(-3.01 - -1.46) | 0.35<br>(-0.06 - 0.8) | 21 | 13 | 272 | 558 |
| Secondary Yng | Scavenger | BL | -1.34<br>(-2.74 - 0.06) | 0.32<br>(-0.14 - 0.79) | 2 | 1 | 25 | 50 |
| Secondary Yng | Aquatic predator | BL | -2.28<br>(-3.09 - -1.48) | 0.3<br>(-0.18 - 0.76) | 20 | 10 | 189 | 457 |
| Secondary Indeter | Nectarivore | BL | -1.15<br>(-1.93 - -0.28) | 0.05<br>(-0.46 - 0.49) | 8 | 2 | 11 | 44 |
| Secondary Indeter | Frugivore | BL | -0.94<br>(-1.71 - -0.2) | 0.08<br>(-0.33 - 0.48) | 16 | 5 | 69 | 185 |
| Secondary Indeter | Granivore | BL | -1.54<br>(-2.2 - -0.83) | 0.11<br>(-0.3 - 0.56) | 20 | 7 | 162 | 979 |
| Secondary Indeter | Herbivore terrestrial | BL | -1.59<br>(-2.73 - -0.44) | 0.07<br>(-0.45 - 0.56) | 1 | 1 | 77 | 77 |
| Secondary Indeter | Herbivore aquatic | BL | -2.09<br>(-3.22 - -1.09) | 0.13<br>(-0.35 - 0.85) | 2 | 2 | 80 | 80 |
| Secondary Indeter | Omnivore | BL | -1.13<br>(-1.65 - -0.57) | 0.04<br>(-0.31 - 0.34) | 66 | 15 | 301 | 2,213 |
| Secondary Indeter | Invertivore | BL | -1.51<br>(-1.99 - -1.04) | 0.08<br>(-0.2 - 0.38) | 130 | 15 | 335 | 3,488 |
| Secondary Indeter | Vertivore | BL | -2.35<br>(-3.18 - -1.56) | 0.11<br>(-0.35 - 0.56) | 20 | 14 | 298 | 588 |
| Secondary Indeter | Aquatic predator | BL | -2.38<br>(-3.21 - -1.56) | 0.06<br>(-0.41 - 0.53) | 13 | 6 | 62 | 122 |
| Cropland | Nectarivore | BL | -1.56<br>(-2.38 - -0.8) | 0.63<br>(0.24 - 1.16) | 14 | 2 | 64 | 439 |
| Cropland | Frugivore | BL | -1.18<br>(-1.83 - -0.53) | 0.61<br>(0.25 - 1) | 30 | 6 | 183 | 1,029 |
| Cropland | Granivore | BL | -1.36<br>(-2.01 - -0.61) | 0.63<br>(0.23 - 1.06) | 28 | 14 | 638 | 2,125 |
| Cropland | Herbivore terrestrial | BL | -1.68<br>(-2.83 - -0.58) | 0.6<br>(0.1 - 1.1) | 2 | 2 | 90 | 90 |
| Cropland | Herbivore aquatic | BL | -2.18<br>(-3.22 - -1.22) | 0.65<br>(0.2 - 1.37) | 7 | 6 | 287 | 493 |
| Cropland | Omnivore | BL | -1.21<br>(-1.72 - -0.74) | 0.63<br>(0.34 - 0.99) | 80 | 18 | 881 | 3,974 |
| Cropland | Invertivore | BL | -1.75<br>(-2.21 - -1.32) | 0.5<br>(0.22 - 0.76) | 179 | 18 | 884 | 8,672 |
| Cropland | Vertivore | BL | -2.44<br>(-3.24 - -1.68) | 0.66<br>(0.27 - 1.15) | 19 | 14 | 749 | 1,993 |
| Cropland | Aquatic predator | BL | -2.52<br>(-3.25 - -1.77) | 0.55<br>(0.06 - 0.97) | 32 | 10 | 543 | 1,952 |
| Pasture | Nectarivore | BL | -1.58<br>(-2.3 - -0.79) | 0.13<br>(-0.31 - 0.58) | 19 | 3 | 36 | 228 |
| Pasture | Frugivore | BL | -1.47<br>(-2.11 - -0.79) | 0.21<br>(-0.14 - 0.69) | 29 | 4 | 55 | 387 |
| Pasture | Granivore | BL | -1.73<br>(-2.38 - -1.08) | 0.12<br>(-0.28 - 0.52) | 30 | 13 | 649 | 1,816 |
| Pasture | Herbivore<br>terrestrial | BL | -1.98<br>(-3.06 - -0.85) | 0.13<br>(-0.36 - 0.62) | 3 | 3 | 72 | 72 |
| Pasture | Herbivore<br>aquatic | BL | -2.39<br>(-3.39 - -1.45) | 0.19<br>(-0.25 - 0.88) | 6 | 4 | 259 | 523 |
| Pasture | Omnivore | BL | -1.53<br>(-2.01 - -1.05) | 0.08<br>(-0.21 - 0.35) | 94 | 16 | 685 | 3,285 |
| Pasture | Invertivore | BL | -1.88<br>(-2.31 - -1.43) | 0.14<br>(-0.13 - 0.42) | 197 | 15 | 671 | 6,583 |
| Pasture | Vertivore | BL | -2.46<br>(-3.19 - -1.72) | 0.14<br>(-0.27 - 0.59) | 21 | 14 | 674 | 1,681 |
| Pasture | Aquatic<br>predator | BL | -2.61<br>(-3.27 - -1.92) | 0.1<br>(-0.38 - 0.53) | 35 | 9 | 619 | 2,596 |
| Plantation<br>Forest | Nectarivore | BL | -0.97<br>(-1.71 - -0.19) | 0.32<br>(-0.08 - 0.78) | 23 | 7 | 35 | 296 |
| Plantation<br>Forest | Frugivore | BL | -1.09<br>(-1.7 - -0.46) | 0.35<br>(0.05 - 0.74) | 45 | 14 | 303 | 1,080 |
| Plantation<br>Forest | Granivore | BL | -1.26<br>(-1.94 - -0.62) | 0.32<br>(-0.05 - 0.71) | 31 | 17 | 352 | 1,498 |
| Plantation<br>Forest | Herbivore<br>terrestrial | BL | -1.49<br>(-2.63 - -0.41) | 0.3<br>(-0.19 - 0.84) | 2 | 2 | 81 | 81 |
| Plantation<br>Forest | Herbivore<br>aquatic | BL | -2.12<br>(-3.21 - -1.11) | 0.37<br>(-0.05 - 1.08) | 4 | 3 | 122 | 204 |
| Plantation<br>Forest | Omnivore | BL | -1.29<br>(-1.78 - -0.82) | 0.31<br>(0.04 - 0.63) | 99 | 27 | 626 | 3,898 |
| Plantation<br>Forest | Invertivore | BL | -1.39<br>(-1.81 - -0.96) | 0.22<br>(-0.03 - 0.44) | 240 | 28 | 676 | 6,979 |
| Plantation<br>Forest | Vertivore | BL | -2.19<br>(-2.97 - -1.46) | 0.33<br>(-0.05 - 0.75) | 25 | 17 | 410 | 640 |
| Plantation<br>Forest | Scavenger | BL | -1.51<br>(-2.91 - -0.13) | 0.34<br>(-0.1 - 0.89) | 1 | 1 | 1 | 1 |
| Plantation<br>Forest | Aquatic<br>predator | BL | -2.23<br>(-2.94 - -1.47) | 0.24<br>(-0.22 - 0.64) | 30 | 11 | 244 | 745 |
| Urban | Nectarivore | BL | -1.62<br>(-2.51 - -0.59) | 0.32<br>(-0.16 - 0.84) | 3 | 2 | 80 | 120 |
| Urban | Frugivore | BL | -1.6<br>(-2.43 - -0.74) | 0.4<br>(-0.09 - 0.96) | 2 | 2 | 127 | 127 |
| Urban | Granivore | BL | -2.05<br>(-2.86 - -1.25) | 0.43<br>(-0.08 - 1.05) | 13 | 5 | 171 | 903 |
| Urban | Herbivore<br>terrestrial | BL | -2.13<br>(-3.35 - -0.96) | 0.35<br>(-0.22 - 0.93) | 1 | 1 | 87 | 87 |
| Urban | Herbivore<br>aquatic | BL | -2.5<br>(-3.61 - -1.38) | 0.43<br>(-0.13 - 1.16) | 1 | 1 | 87 | 87 |
| Urban | Omnivore | BL | -1.56<br>(-2.26 - -0.87) | 0.32<br>(-0.15 - 0.77) | 23 | 4 | 170 | 1,547 |
| Urban | Invertivore | BL | -2.35<br>(-2.98 - -1.76) | 0.29<br>(-0.17 - 0.75) | 49 | 4 | 170 | 3,656 |
| Urban | Vertivore | BL | -2.75<br>(-3.62 - -1.79) | 0.4<br>(-0.12 - 0.96) | 7 | 1 | 3 | 21 |
| Urban | Aquatic predator | BL | -2.9<br>(-3.82 - -1.92) | 0.33<br>(-0.25 - 0.9) | 1 | 1 | 3 | 3 |

**Table S6.** The number of distinct values for each hierarchical of the Bayesian hierarchical logistic regression model. Values are given for data used to fit models using global abundance estimates sourced from Callagahan *et al*. (*19*) and BirdLife International (*24*).

| **Hierarchical level** | **‘n’ Callaghan** | **‘n’ Bird-Life** |
| --- | --- | --- |
| *i* (detections / non-detections) | 395,305 | 119,401 |
| *l* (land-cover types) | 9 | 9 |
| *k* (species) | 3,146 | 798 |
| *tn* (trophic niches) | 10 | 10 |
| *s* (PREDICTS studies) | 102 | 95 |
| *j* (PREDICTS study sites) | 5,454 | 5,262 |

